# Adjusting for Common Variant Polygenic Scores Improves Yield in Rare Variant Association Analyses

**DOI:** 10.1101/2021.10.18.464866

**Authors:** Sean J. Jurgens, James P. Pirruccello, Seung Hoan Choi, Valerie N. Morrill, Mark Chaffin, Steven A. Lubitz, Kathryn L. Lunetta, Patrick T. Ellinor

## Abstract

With the emergence of large-scale sequencing data, methods for improving power in rare variant analyses (RVAT) are needed. Here, we show that adjusting for common variant polygenic scores improves the yield in gene-based RVAT across 65 quantitative traits in the UK Biobank (up to 20% increase at α=2.6×10^−6^), without a marked increase in false-positive rates or genomic inflation. Our results illustrate how adjusting for common variant effects can aid in rare variant association discovery.

---

In recent years, large-scale biorepositories have seen an explosion in available high-depth sequencing data^1,2^, and investigators have increasingly leveraged gene-based tests to identify rare variants contributing to human phenotypic variability^3–5^. An important direction in the genetics field is to identify methods for improved power in rare variant association analyses (RVAT). Many quantitative traits have considerable heritability from common variants^6^. We therefore hypothesized that accounting for the effects attributable to common variants would improve power in RVAT. We leveraged the UK Biobank dataset, which contains imputed data on nearly 500,000 individuals^7^ as well as exome sequencing for over 200,000 individuals^2^. We show that adjusting for common variant effects, summarized in polygenic scores (PRS), improves the yield in gene-based RVAT across 65 quantitative traits.

We first performed genome-wide association analyses (GWAS) for common variants (MAF≥1%) across 65 quantitative traits (**Supplementary Table 1**). We performed three types of GWAS, namely an out-of-sample GWAS within European samples who were not included in the exome sequencing subset (N=230k), an in-sample GWAS within European samples who were also exome sequenced (N=190k), and a ‘total’ GWAS within all European UK Biobank participants (N=460k) (**Supplementary Figure 1**). All traits had multiple independent genome-wide significant (*P*<5×10^−8^) common variant hits (**Supplementary Figure 2A**, **Supplementary Table 2**).

Using the GWAS summary statistics, we then constructed PRS based on two methods, namely ‘lead-SNP’ PRS (*P*<5×10^−8^ and r^2^<0.001), and genome-wide PRS using PRScs-auto^8^ (**Methods**). Thus, we analyzed six PRS per trait: PRS_lead-SNP (out-sample)_, PRS_CS (out-sample)_, PRS_lead-SNP (in-sample)_, PRS_CS (in-sample)_, PRS_lead-SNP (total)_ and PRS_CS (total)_. All types of PRS explained variance for their respective traits (**Supplementary Figure 2B, Supplementary Table 2**).

We then performed exome-wide gene-based collapsing RVAT within the exome sequenced samples, focusing on ultra-rare loss-of-function (LOF) and missense variants with MAC<40 (**Methods**). We ran RVAT models with no PRS included, as well as RVAT models adjusting for each type of PRS.

All six PRS-adjusted models showed higher numbers of RVAT gene-phenotype associations at various significance cutoffs, compared to the model without a PRS (**Figure 1A**, **Supplementary Figure 3**, **Supplementary Tables 3-4**). PRS_CS (out-sample)_ generally yielded more total associations than PRS_lead-SNP (out-sample)_. The PRS_CS (out-sample)_ model yielded 13.3% and 19.7% more significant associations at Bonferroni-corrected significance (α=7.2×10^−8^; 170 vs 150 associations), and conventional exome-wide significance (α=2.6×10^−6^; 261 vs 218 associations), respectively. PRS_lead-SNP (in-sample)_ performed similarly to PRS_lead-SNP (out-sample)_, while PRS_CS (in-sample)_ generally performed the least well (**Figure 1A**).

**Figure 1:**
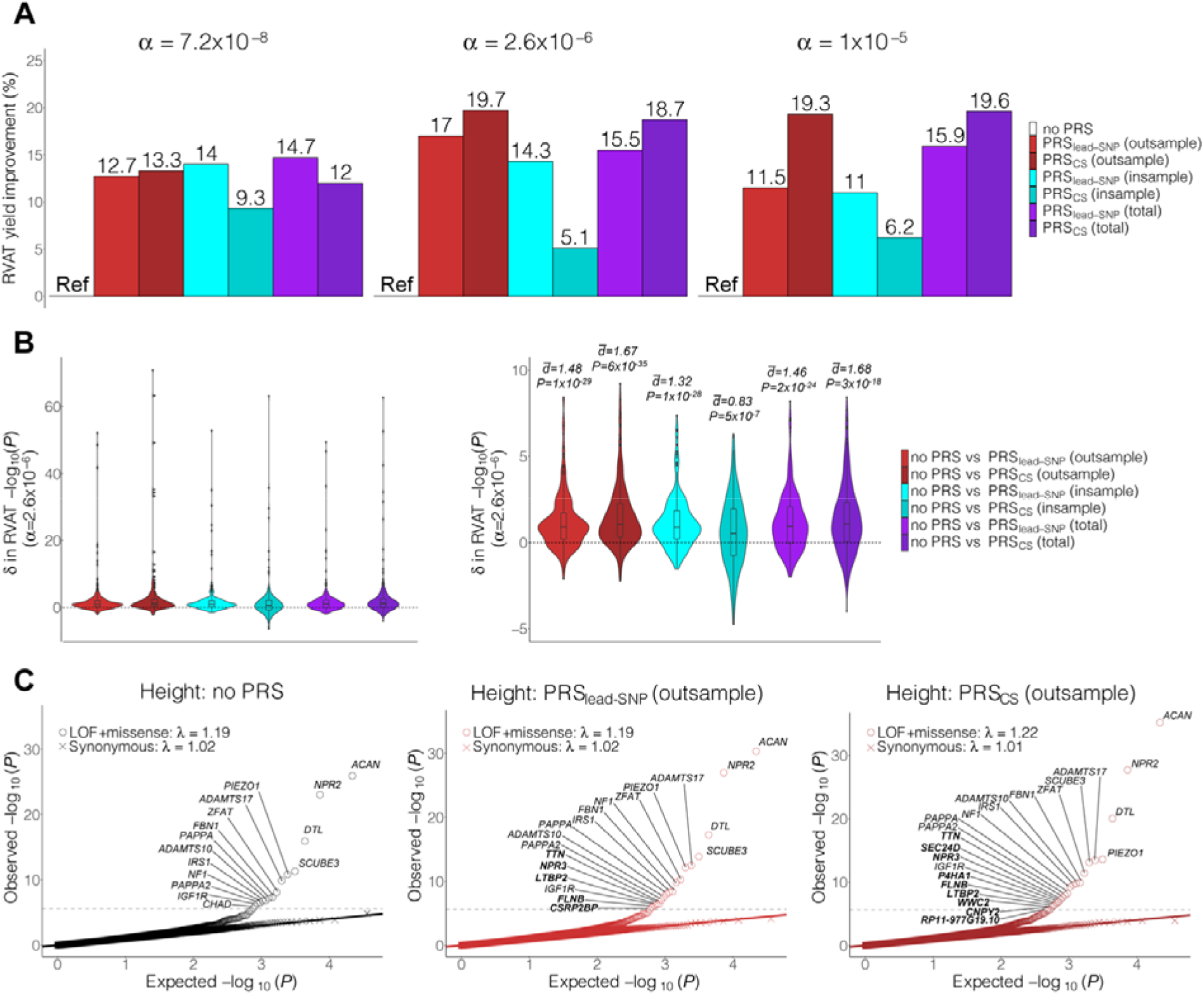
PRS adjustment improves discovery yield in analysis of rare deleterious variants. **Panel A**: Bar charts for the improvement in deleterious RVAT yield after PRS-adjustment at different alpha levels, expressed in percentage relative to the no PRS model. **Panel B**: Violin plots for the difference (δ) in significance of tests from deleterious RVAT, comparing models with PRS vs models without PRS. Here, the δ in *P*-values (on the −log_10_ scale) are displayed for tests reaching *P*<2.6×10^−6^ (Methods). The d values plotted above the violins are derived from two-sided paired T-tests (after removing outliers), while the *P*-values are derived from two-sided paired Wilcoxon signed rank tests. The left plot shows all results, while the right plot is capped at y=10 for clarity. Boxplots: center line, median; box limits, upper and lower quartiles; whiskers, 1.5× interquartile range; points, outliers. **Panel C**: Quantile-quantile plots for PRS-adjusted RVAT of the phenotype height. The left plot shows expected vs observed *P*-values for the model with no PRS-adjustment, while the second and third plots show results for PRS_leadSNP (out-sample)_ and PRS_CS (out-sample)_, respectively. Exome-wide significant genes are annotated with gene names; genes highlighted in bold were only identified after PRS-adjustment.

At various significance thresholds, PRS-adjusted models significantly improved the *P*-values for top gene-phenotype associations, as compared to the model without PRS (**Supplementary Figure 4** and **Supplementary Table 5**). For example, the PRS_CS (out-sample)_ adjusted model was associated with significantly higher −log10(*P*) values, for associations reaching conventional exome-wide significance (*P*=6×10^−35^, paired Wilcoxon signed-rank test) (**Figure 1B**; **Supplementary Table 5**).

Many of the gene-phenotype associations that became significant after PRS-adjustment were biologically-plausible findings (**Supplementary Note**, **Supplementary Table 6**). As an example, for the phenotype height such associations included *NPR3*, *LTBP2*, *P4HA1*, *FLNB*, *SEC24D* and *TTN* (**Figure 1C**, **Supplementary Note, Supplementary Figure 5**).

We found that h^2^_SNP_ was significantly associated with the per-trait improvement in number of significant associations after PRS-adjustment, particularly for PRS_CS_ models (**Supplementary Figure 6**). Similarly, PRS R^2^ was a significant positive predictor for the per-trait change in association yield (**Supplementary Figure 7**).

To assess genomic inflation and false-positive rates, we then performed exome-wide RVAT analyzing synonymous variants with MAC<40. At liberal α cutoffs, we observed association rates that were marginally higher or equivalent to the expectation under the null (**Figure 2A**, **Supplementary Figure 3**, **Supplementary Table 7**). At Bonferroni-corrected significance (α=4.3×10^−8^), we observed more hits than expected under the null (**Supplementary Tables 8-9**). We found that all these associations involved *IGLL5* and white blood cell traits (**Supplementary Table 10**), possibly reflecting true association^9^. After removing *IGLL5* from the analysis, synonymous association rates were well controlled at stringent α values (**Supplementary Table 7**).

**Figure 2:**
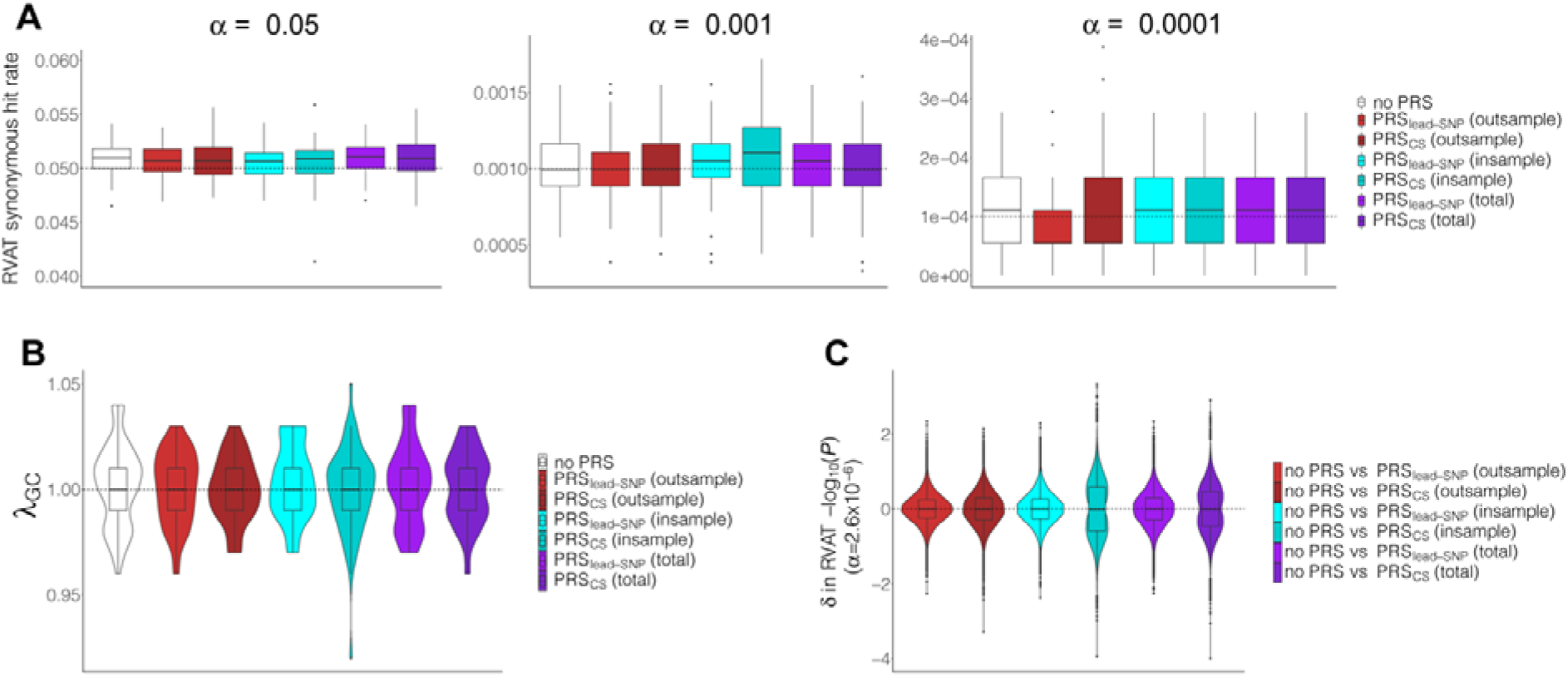
PRS adjustment does not increase false-positive rates or genomic inflation in the analysis of rare synonymous variants. **Panel A**: Boxplots for per-trait association rate from synonymous RVAT at different alpha levels across the 65 traits. Per trait, a median of 18,060 genes were analyzed. **Panel B**: Violin plots for genomic inflation factors for exome-wide RVAT of synonymous variants across the 65 traits. **Panel C**: Violin plots for difference (δ) in significance of tests from synonymous variant RVAT, comparing models with PRS vs models without PRS. Here, the δ in *P*-values (on the −log10 scale) are displayed for tests reaching *P*<0.05 (Methods). The d values plotted above the violins is derived from two-sided paired T-tests (after removing outliers) while the *P*-values are derived from two-sided paired Wilcoxon signed-rank tests. Boxplots: center line, median; box limits, upper and lower quartiles; whiskers, 1.5× interquartile range; points, outliers.

Importantly we did not observe a clear pattern where synonymous association rates were strongly increased after PRS adjustment. Using paired Wilcoxon signed-rank tests, we found no significant increase in −log10(*P*) values for the synonymous RVAT at various α levels, across the different types of PRS adjusted models (*P*>0.05 for all tests by paired Wilcoxon rank test; **Supplementary Figure 4** and **Supplementary Table 11**). For example, at the α=0.05 level, estimated differences between models with PRS vs without PRS centered around 0 (**Figure 2C**, **Supplementary Table 11**).

We then assessed inflation factors, using per-trait λ_GC_ values (**Supplementary Tables 12-13**). In synonymous RVAT, per-trait λ_GC_ values did not increase after PRS adjustment across PRS types (**Supplementary Figure 8**, **Supplementary Table 13**). All per-trait synonymous λ_GC_ values were within acceptable limits at λ_GC_<1.05 (**Figure 2B**), and test statistics were not inflated visually (**Supplementary Figure 9**).

In conclusion, we find that adjustment for common variant PRS can improve the yield in gene-based RVAT, without markedly increasing false-positive rates and genomic inflation. The observed power increase likely reflects true biological variance being absorbed by PRS. Indeed, the genome-wide PRS_CS_ performed better than PRS_lead-SNP_ for out-of-sample GWAS data. Further, not all traits had equal benefit from PRS adjustment (sometimes having decreased yield), with SNP-heritability and PRS R^2^ being strong positive predictors of yield improvement. We note that for in-sample GWAS, PRS_CS_ did not perform as well as other PRS, likely owing to overfitting of this genome-wide model. We therefore recommend using large out-of-sample GWAS data when available, or using simple PRS models when independent GWAS data is not available.

Our analysis was focused on burden testing of ultra-rare variants for quantitative traits. While our approach may not be optimal for low-frequency variants, it is useful for ultra-rare variants, which are of particular interest in contemporary sequencing studies. Furthermore, additional studies will be needed to evaluate PRS-adjustments for kernel-based RVAT methods and for binary traits.

In sum, we show how adjusting for common variant effects can aid in rare variant association discovery. Our approach can be applied to enhance discovery yield in future rare variant analyses.

## Data Availability

Summary statistics from the common variant association analyses, summary statistics from the rare variant association analyses, and summary data for polygenic score construction will be made available for download through the Cardiovascular Disease Knowledge Portal upon publication (https://cvd.hugeamp.org). Summary statistics for the tests of the statistical properties of different RVAT models are included in the **Supplementary Tables**. Access to individual level UK Biobank data, both phenotypic and genetic, is available to bona fide researchers through application on the UK Biobank website (https://www.ukbiobank.ac.uk).

## Software and Code Availability

Quality-control of individual level data was performed using Hail version 0.2 (https://hail.is) as well as PLINK version 2.0.a (https://www.cog-genomics.org/plink/2.0/). Variant annotation was performed using VEP version 95 (https://github.com/Ensembl/ensembl-vep). Main common variant association analyses (GWAS) were performed using REGENIE v2.0.2 (https://github.com/rgcgithub/regenie). Genome-wide polygenic scores were computed using PRS-CS (https://github.com/getian107/PRScs; githash: 43128be7fc9ca16ad8b85d8754c538bcfb7ec7b4). Main rare variant association analyses were performed using an adaptation of the R package GENESIS version 2.18 (https://rdrr.io/bioc/GENESIS/man/GENESIS-package.html), which has previously been made available by us through the GitHub repository https://github.com/seanjosephjurgens/UKBB_200KWES_CVD. Analyses were run within R version 4.0 (https://www.r-project.org).

## Online Methods

### Study population

The UK Biobank is a large population-based prospective cohort study from the United Kingdom with rich phenotypic and genetic data on 500,000 individuals aged 40-69 at enrollment^10^. Available genetic data currently includes genome-wide imputed data for almost all participants^7^, as well as whole exome sequencing data on approximately 200,000 individuals^2^. The UK Biobank resource was approved by the UK Biobank Research Ethics Committee and all participants provided written informed consent to participate. Use of UK Biobank data was performed under application number 17488 and was approved by the local Massachusetts General Hospital Institutional Review Board.

### Phenotypes

In the present study, we analyzed 65 quantitative traits, including anthropometric traits, metabolic blood markers, blood pressure traits, and a variety of blood count traits. Details and the number of samples for each trait per analysis are presented in **Supplementary Table 1**. All raw phenotypes were adjusted for lipid-lowering medication use (**Supplementary Note**), and were subsequently rank-based inverse normalized to ensure normality before analyses.

### Genetic datasets

We utilized both genome-wide imputed data and whole exome sequencing data in the present study. Specifically, all common variant analyses were performed using genome-wide imputed data^7^. Briefly, genotyping was performed using Affymetrix UK Biobank Axiom (450,000 samples) and Affymetrix UK BiLEVE axiom (50,000 samples) arrays. Subsequently, the genetic data were imputed to the Haplotype Reference Consortium panel and UK10K + 1000 Genomes panels. We removed any samples that had withdrawn their consent, samples that were outliers for heterozygosity or missingness, individuals with putative sex chromosome aneuploidy, and individuals with a mismatch between self-reported and genetically inferred sex. We then removed all individuals who were determined to not be of homogeneous European ancestry (**Supplementary Note**). To ensure we analyzed only high-quality common imputed variants, we removed imputed variants with minor allele frequency (MAF) <1% and INFO <0.3.

For all rare variant analyses, we utilized the whole exome sequencing data, which were available for 200,642 individuals^2^. The revised version of the IDT xGen Exome Research Panel v1.0 was used to capture exomes with over 20X coverage at 95% of sites. Variants were subsequently called per-sample using DeepVariant and combined using GLNexus^11^. We utilized the quality-control procedures described previously in Jurgens et al. (under review^5^). In short, we set low-quality genotypes to missing, after which we removed variants based on call rate (<90%), Hardy-Weinberg equilibrium test (*P* < 1×10^−15^), presence in low-complexity regions, and minor allele count (≥1). Sample-level quality-control consisted of removal of samples that had withdrawn their consent, were duplicates, had a mismatch between sequencing and genotyping array data, had a mismatch between genetically inferred and self-reported sex, had low call rates or were outliers for a number of additional metrics (Jurgens et al., under review^5^). We finally restricted the exome cohort to individuals who also had imputed data available and were of European ancestry, leaving 188,062 samples.

### Common variant association analyses

We first performed three genome-wide association analyses (GWAS) for each included trait using genome-wide imputed data (**Supplementary Figure 1**). These included an out-of-sample GWAS within European samples who were independent of the exome cohort (not included in the exome cohort and unrelated to the exome cohort); an in-sample GWAS within the exome sequenced samples; and a total GWAS including all European individuals with imputed data. To perform the GWAS, we used linear whole-genome ridge regression models implemented in REGENIE^12^, adjusting for sex, age, age^2^, genotyping array and ancestral principal components 1 through 20. REGENIE produces results similar to linear mixed models in the presence of genetic relatedness^12^.

### Polygenic score derivation

Using each of the GWAS summary results, we constructed polygenic scores (PRS) for each trait based on two differing methods^13^. We first constructed ‘lead SNP’ PRSs based only on independent (r^2^<0.001) genome-wide significant (*P*<5×10^−8^) variants. We also used PRS-CS-auto^8^ to construct genome-wide PRSs including millions of genetic variants (restricting to ∼1.1 million HapMap variants). In brief, PRS-CS-auto applies a Bayesian regression framework to identify posterior variant effect sizes based on a continuous shrinkage prior, which is directly learnt from the data^8^. For both methods, the European ancestry subset of the UK Biobank dataset was used as a linkage-disequilibrium reference panel. In sum, two PRS were constructed for each trait based on out-sample GWAS data (PRS_leadSNP [out-sample]_ and PRS_CS [out-sample]_), two PRS were constructed based on in-sample GWAS data (PRS_leadSNP [in-sample]_ and PRS_CS [in-sample]_) and two PRS were constructed based on total GWAS data (PRS_leadSNP [total]_ and PRS_CS [total]_).

### Variance explained by PRS

We calculated the phenotypic variance explained by each PRS for each trait in the nullmodel. We did this by running ordinary linear regression for each trait among the unrelated subset of individuals with exome sequencing data, adjusting for the same fixed effects as described above for the rare variant analysis. R^2^ values were extracted from the model without PRS and from models with PRS added as a covariate. The variance explained by PRS for a given trait was defined as the improvement in R^2^ in the model with PRS as compared to the model with no PRS.

### Rare variant association analyses

We used the whole exome sequencing data to run gene-based rare variant collapsing tests across the exome for each trait. We grouped and analyzed loss-of-function (LOF) and predicted-deleterious missense variants per gene (**Supplementary Note**). To ensure no linkage between common and rare variants, we only included variants with minor allele count (MAC) ≤40, which also had MAF<0.1% in each continental population in gnomAD version 2 exomes^14^. We utilized linear mixed models implemented in GENESIS^15^, adjusting for sex, age, age^2^, genotyping array, sequencing batch, ancestral principal components 1 through 20, and a sparse kinship matrix (Jurgens et al., under review^5^). We subsequently repeated these analyses for each of the PRS, by adding the PRS to the model as an additional fixed-effect covariate. In cases where fitting of the mixed model failed, we reran models within unrelated individuals (**Supplementary Table 1**). Sample sizes for the rare variant analyses ranged from N=142,709 to N=187,890 (**Supplementary Table 1**). Only results for tests with ≥20 rare variant carriers were kept.

### Assessment of rare variant discovery yield

We then evaluated the rare variant discovery power for models without PRS and those adjusted for PRS. We calculated the yield in number of gene associations for each model across all traits at various significance thresholds, including Bonferroni-corrected significance at α = 0.05/ (65 traits × ∼10,743 genes) = 7.2×10^−8^, and at conventional exome-wide significance at α = 2.6×10^−6^. We then tested whether the addition of the PRS improved the significance of gene-phenotype associations. We used two-sided paired Wilcoxon signed rank tests to assess the improvement in −log_10_(*P*) values between two models, including gene-phenotype associations at various significance cutoffs (7.2×10^−8^, 2.6×10^−6^, 1×10^−5^, 1×10^−4^, 1×10^−3^, 0.05). For a given comparison between two models, we included any gene-phenotype pair reaching the cutoff in either model. To quantify the difference, d□, in −log_10_(*P*) values, we repeated this analysis using paired T-tests. For paired T-tests, we removed any gene-phenotype pair for which the difference between both models fell outside of 4 standard deviations from the mean of differences. The significance threshold was determined at α = 0.05 / (6 cutoffs × 6 model comparisons) = 0.0014.

### Associations between trait heritability and PRS variance explained with yield improvement

We then assessed whether the improvement in RVAT associations after PRS-adjustment was associated with trait heritability or the variance explained by PRS. We used Linkage-Disequilibrium Score Regression^16^ to estimate SNP-heritability (h^2^_SNP_) for each of the 65 traits, using the total sample GWAS results and using the *baselineLD_2.2* file from the LDSC software as the linkage-disequilibrium reference. We then used ordinary linear regression to regress the change in number of trait RVAT associations on the estimated h^2^_SNP_. Similary, we used linear regression to regress the change in number of trait RVAT associations on the R^2^ of the PRS for its respective traits.

### Assessment of false-positive rate using rare synonymous variation

To assess the false-positive error rate of our approach, we analyzed rare synonymous variation. Synonymous variants are generally not expected to affect the amino acid sequence encoded by genes, and therefore are strongly depleted of true genetic effects^17^. We grouped rare synonymous variants (MAC≤40 and MAF<0.1% in each continental population in gnomAD exomes) and ran exome-wide gene-based collapsing tests using GENESIS. Only results for tests with ≥20 rare variant carriers were kept. The significant association rate for synonymous variants was determined at various significance cutoffs for each model: α = 4.3×10^−8^ (Bonferroni-corrected), 2.6×10^−6^, 1×10^−5^, 1×10^−4^, 1×10^−3^, and 0.05. As described for the deleterious variants above, we further utilized paired Wilcoxon rank tests and paired T-tests to evaluate the changes in −log_10_(*P*) values at different significance levels.

### Assessment of exome-wide inflation

Exome-wide test statistics were plotted in quantile-quantile (QQ) plots to visually assess inflation per trait, per model, per variant mask. Exome-wide inflation was further quantified using λ-values, defined as the empirical χ^2^ statistic at the median divided by the expected χ^2^ statistic at the median under the null. To assess whether λ-values differed between models without PRS and those adjusted for various PRS across the 65 traits, we utilized two-sided paired Wilcoxon rank tests and paired T-tests.

