## Supplementary Information for "Adjusting for Common Variant Polygenic Scores Improves Yield in Rare Variant Association Analyses"

Sean J. Jurgens^1,2,*^, James P. Pirruccello^1,*^, Seung Hoan Choi^1^, Valerie N. Morrill^1^, Mark Chaffin^1^, Steven A. Lubitz^1,5,6^, Kathryn L. Lunetta^3,4^ & Patrick T. Ellinor^1,5,6^

1. Cardiovascular Disease Initiative, The Broad Institute of MIT and Harvard, Cambridge, MA, USA

2. Department of Experimental Cardiology, Amsterdam UMC, University of Amsterdam, Netherlands, NL

3. NHLBI and Boston University’s Framingham Heart Study, Framingham, MA, USA

4. Department of Biostatistics, Boston University School of Public Health, Boston, MA, USA

5. Demoulas Center for Cardiac Arrhythmias, Massachusetts General Hospital, Boston, MA, USA

6. Cardiovascular Research Center, Massachusetts General Hospital, Boston, MA, USA

^*^ Contributed equally to this work.

| **Items** | | **Pages** |
| --- | --- | --- |
| **Supplementary Note: 1. Supplementary Methods** | | |
|  | **Lipid-lowering medication adjustment** | 4 |
|  | **Identifying individuals of homogeneous European ancestry** | 4 |
|  | **Variant included in the rare deleterious variant analyses** | 4-5 |
| **Supplementary Note: 2. Supplementary Results** | | |
|  | **Biologically plausible findings among genes becoming significant after PRS-adjustment** | 5-6 |
| **Supplemental Tables** | | |
|  | **Supplementary Table 1: Characteristics of phenotypes and performed analyses** | Excel file |
|  | **Supplementary Table 2: Number of variants included in the polygenic scores and variance explained in the nullmodels** | Excel file |
|  | **Supplementary Table 3: Number of significant gene associations and inflation factors for analysis of rare loss-of-function and deleterious missense variants** | Excel file |
|  | **Supplementary Table 4: Number of significant gene associations for rare loss-of-function and deleterious missense analysis at different significance cutoffs** | Excel file |
|  | **Supplementary Table 5: Improvement in -log_10_(*P*) values after PRS adjustment in the analysis of LOF and missense variants, at different cutoffs** | Excel file |
|  | **Supplementary Table 6: Significant gene-phenotype associations from LOF and missense analysis at exome-wide significance** | Excel file |
|  | **Supplementary Table 7: False-positive rate based on synonymous variation at different alpha values** | Excel file |
|  | **Supplementary Table 8: Number of significant gene associations and inflation factors for analysis of rare synonymous variants** | Excel file |
|  | **Supplementary Table 9: Number of significant gene associations for rare synonymous variant analysis at different significance cutoffs** | Excel file |
|  | **Supplementary Table 10:** **Significant gene-phenotype associations from synonymous analysis at exome-wide significance significance** | Excel file |
|  | **Supplementary Table 11:** **Improvement in -log_10_(*P*) values after PRS adjustment in the analysis of synonymous variants, at different cutoffs** | Excel file |
|  | **Supplementary Table 12:** **Change in λ values after PRS adjustment in LOF and missense analysis** | Excel file |
|  | **Supplementary Table 13:** **Change in λ values after PRS adjustment in synonymous analysis** | Excel file |
| **Supplemental Figures and Figure Legend** | | |
|  | **Supplementary Figure 1. Analysis Flowchart** | 7 |
|  | **Supplementary Figure 2. Number of significant lead variants from common variant GWAS and variance explained by subsequently derived PRS across the 65 traits** | 8 |
|  | **Supplementary Figure 3.** **Bar plots for RVAT yield at different significance cutoffs** | 9-10 |
|  | **Supplementary Figure 4.** **Violin plots for difference in P-values from models with PRS compared to models without PRS at different significance thresholds.** | 11-13 |
|  | **Supplementary Figure 5: Quantile-quantile plots for the phenotype height with exome-wide significant genes annotated.** | 14-15 |
|  | **Supplementary Figure 6.** **Correlation between SNP-heritability and the improvement in number of significant rare variant associations after PRS adjustment across the 65 traits.** | 16 |
|  | **Supplementary Figure 7: Correlation between PRS variance explained and the change in the number of significant rare variant associations after PRS adjustment across the 65 traits.** | 17 |
|  | **Supplementary Figure 8: Paired box plots for comparison of genomic inflation factors between models without PRS and models with PRS** | 18-19 |
|  | **Supplementary Figure 9: Quantile-quantile plots for exome-wide rare variant analysis of deleterious variants and synonymous variants by model.** | 20-41 |
| **Supplementary References** | | 42 |

**Supplementary Note**

***Supplementary Methods***

**Lipid-lowering medication adjustment**

For all traits studied in this manuscript, the UK Biobank reported measurements at two separate visits for a subset of participants. We identified 20,157 individuals with measurements from at least two visits, of whom 15,952 individuals were not on lipid-lowering therapy at the time of the first visit. Of these participants, we identified 2,292 individuals who did not report taking a lipid lowering therapy (Anatomical Therapeutic Chemical [ATC] code beginning with C10) at the initial visit, but did report taking one at the second visit. The ATC codes were mapped from UK Biobank drug names by Wu *et al*^1^. Separately for each trait, we constructed a linear model computing the quantitative trait’s value at the second visit as a function of the trait’s value at the first visit, adjusted for the age at the time of the second visit, sex, and a dichotomous variable representing whether the participant had initiated a lipid-lowering therapy between the first and second visit (true for 2,292 individuals and false for 13,660). We treated the effect size of the dichotomous variable as an additive constant for that trait. We used these additive constants in the entire UK Biobank population to adjust raw trait values. As an example, if lipid-lowering therapies were found to be associated with a reduction in LDL cholesterol by 0.94 mmol/liter, we added 0.94 mmol/liter to the LDL cholesterol values for all individuals taking lipid-lowering therapies.

**Identifying individuals of homogeneous European ancestry**

We restricted to individuals of genetically-determined European ancestry in all of our analyses. To this end, we utilized ADMIXTURE^2^ to compute likelihoods for 5 major continental super-populations (European, African, South Asian, East Asian and Admixed American ancestries). Models were trained using 2,504 samples with whole genome sequencing data from the 1000 Genomes project, for whom global ancestries were known^3^. Variants from the 1000 Genomes dataset were kept if they had MAF>=1% in at least one of the major super-populations; had overall missingness <2%; were non-ambiguous; and had Hardy-Weinberg equilibrium test *P*>0.0001 in each of the major super-populations. Variants were then restricted to those also found with missingness <2% in the UK Biobank genotyping array data. Finally, variants were consecutively pruned within each major continental population from the 1000 Genomes dataset, utilizing PLINK2^4^ with the command *--indep-pairwise 50 5 0.2*. These procedures left 87,398 variants which were used to train ADMIXTURE models within the 1000 Genomes samples, and for subsequent likelihood estimation in the UK Biobank samples with genotyping array data. Individuals from the UK Biobank with >80% probability for European ancestry were considered of homogenous European ancestry, leaving 459,589 individuals.

**Variant included in the rare deleterious variant analyses**

The protein consequences of variants were interrogated using dbNSFP^5^ (version 4.1a) and the Loss-of-Function Transcript Effect Estimator^6^ (LOFTEE) plug-in implemented in the Variant Effect Predictor^7^ (VEP; version 95) (<https://github.com/konradjk/loftee>). VEP was used to ascertain the most severe consequence of a given variant for each gene transcript. LOFTEE was implemented to identify high-confidence loss-of-function variants (LOF), which include frameshift indels, stopgain variants and splice site disrupting variants. We also removed any LOFs flagged by LOFTEE as dubious, such as LOFs affecting poorly conserved exons and splice variants affecting NAGNAG sites or non-canonical splice regions. Missense variants were annotated using 30 *in silico* prediction tools included in the dbNSFP database.

For missense variants, we calculated a ‘predicted-deleteriousness’ score, based on 30 *in silico* prediction tools. All missense variants were annotated using VEP and the dbNSFP database^5^ (version 4.1a.) The tools from this database included a number of qualitative prediction algorithms (SIFT, SIFT4G, Polyphen2 HDIV, Polyphen2 HVAR, LRT, MutationTaster, FATHMM, PROVEAN, MetaSVM, MetaLR , MCAP, PrimateAI, DEOGEN2, BayesDel addAF, BayesDel noAF, ClinPred, LIST-S2, fathmm-MKL coding, fathmm-XF coding, MutationAssessor, and Aloft) as well as quantitative algorithms (VEST4, REVEL, MutPred, MVP, MPC, DANN, CADD, Eigen, and Eigen-PC). When qualitative prediction tools (except for MutationAssessor and Aloft) indicated "D" for a variant, the variant gained one score from each algorithm. An indicator for a deleterious variant from MutationAssessor was "H" and from Aloft was "R" or "D" with high confidence. For the quantitative algorithms, when the variant indicators were higher than 90% of predicted variants in the entire dataset, the variant gained one score from the quantitative algorithm. Then, if a variant was annotated with more than seven prediction tools (over 20% out of the 30 tools), and the proportion of the deleterious score (total gained score / # none missing prediction tools) was greater than or equal to 0.9, we included the variant in the gene-based analyses of deleterious variants. Final variant masks for analysis of deleterious variants were restricted to high-confidence LOF variants and missense variants with predicted deleterious scores of >=0.9, comparable to the masks used previously^8^.

***Supplementary Results***

**Biologically plausible findings among genes becoming significant after PRS-adjustment**

Among the associations that became exome-wide significant (α=2.6x10^-6^) after adjusting for out-of-sample PRS, many were well-known gene-phenotype associations (**Supplementary Table 6**). These include the association between *CREB3L3*, a gene in which rare variants cause dyslipidemia^9^, with apolipoprotein-a; the association between *PCSK1*, a Mendelian obesity gene^10^, with body-mass-index; the association between *NLRP3*, an inflammasome protein^11^, with c-reactive protein; the associations between *LRP2*, a protein targeted in anti-LRP2 nephropathy^12^, with the renal parameter cystatin-c; the association between *SLC34A3*, a gene in which Mendelian mutations cause renal dysphosphatemias^13^, with cystatin-c; *SLC4A1*, a gene associated with hereditary spherocytosis^14^, with direct and total bilirubin; *TMPRSS6*, a gene in which rare variants cause iron-deficiency anemia^15^, with haematocrit percentage; *RNF10*, a gene overlapping a common variant locus strongly associated with reticulocyte fraction^16^, with highlight scattered reticulocyte count; *EIF2AK1*, of which the protein product has been shown to regulate heme and globin synthesis in model organisms^17^, with immature reticulocyte fraction; *ABCG5*, a Mendelian sitosterolemia gene^18^, with LDL levels; *GAS6*, near which common variants are strongly associated with LDL^19^, with LDL; *CHTF18* and *DNAJC13*, both of which have been associated with mean corpuscular haemaglobin in common variant analyses^16,20^, with mean haemoglobin traits; *SPTA1*, a gene associated with hereditary spherocytosis^14^, with mean corpuscular haemaglobin concentration; *ACTN1*, a gene responsible for ACTN1-related thrombocytopenia^21^, with mean thrombocyte volume; *DIAPH1*, described for autosomal dominant macrothrombocytopenia with hearing loss^22^, with mean thrombocyte volume; *KIAA1109*, near which common variants are associated with platelet volume^16^, with mean platelet volume; *SVEP1*, a gene recently implicated in thrombocyte traits through gene-based RVAT in TOPMed^23^, with mean thrombocyte volume; *ANK1*, *EPB41* and *LCAT*, genes which are known to cause hereditary spherocytosis^14^, hereditary elliptocytosis^24^ and LCAT-deficiency with hemolytic anemia^25^, respectively, were associated with mean sphered cell volume; *RASA3*, a regulator of hematopoiesis with knockout in model organisms causing pancytopenia^26^, with neutrophil percentage; *CASR*, a cause of Mendelian hyperparathyroidism and hypocalciuric hypercalcemia^27^, with phosphate levels; *ITGB3*, a gene known for Glanzmann thrombasthenia^28^, with platelet count; *COL18A1* and *PRC1*, near which common variants are associated with platelet traits^16,29^, with platelet traits; *HBB*, known for beta-thalassemia^30^, with reticulocyte count; *ZNF770*, a gene near which common variants are associated with SHBG levels^31^, with SHBG levels; *ALB*, encoding albumin which is a known binder of bilirubin^32^, with total bilirubin levels; *TNFSF13*, near which common variants are associated with protein levels^33^, with total protein; *PKD1*, an important gene for autosomal dominant polycystic kidney disease^34^, with urate levels; *ABCG2*, encoding transporter important in uric acid excretion^35^, with urate levels; *DHCR7*, encoding an enzyme involved in metabolism of the vitamin-D precursor 7-dehydrocholesterol^36^, with vitamin-D levels; *RC3H1,* a gene described for a human immune dysregulation syndrome^37^, with leukocyte count.

Given the high SNP-heritability of height^38^, we then focused specifically on the RVAT associations for this phenotype. For the phenotype height, several plausible genes became exome-wide significant after PRS adjustment (**Supplementary Table 6**). RVAT associations reaching conventional exome-wide significance (α=2.6x10^-6^) are displayed in **Figure 1C** and **Supplementary Figure 6**. Biologically-plausible associations that became exome-wide significant after PRS adjustment included *NPR3* (associated with connective tissue disorder with increased growth^39^), *LTBP2* (responsible for a syndrome with ocular disease and tall stature^40^), *P4HA1* (a gene for connective tissue disease and myopathy^41^), *FLNB* (associated with Larsen syndrome^42^), *SEC24D* (responsible for syndromic osteogenesis imperfecta^43^), *CNPY2* and *RP11-977G19.10* (which overlap a common variant locus strongly associated with standing height^44^), and *TTN* (associated with skeletal and cardiomyopathies^45^). Across all PRS models, only one gene that was exome-wide significant dropped below significance after PRS adjustment, namely *CHAD* (encoding chondroadherin which is associated with height in common variant analyses^44^).

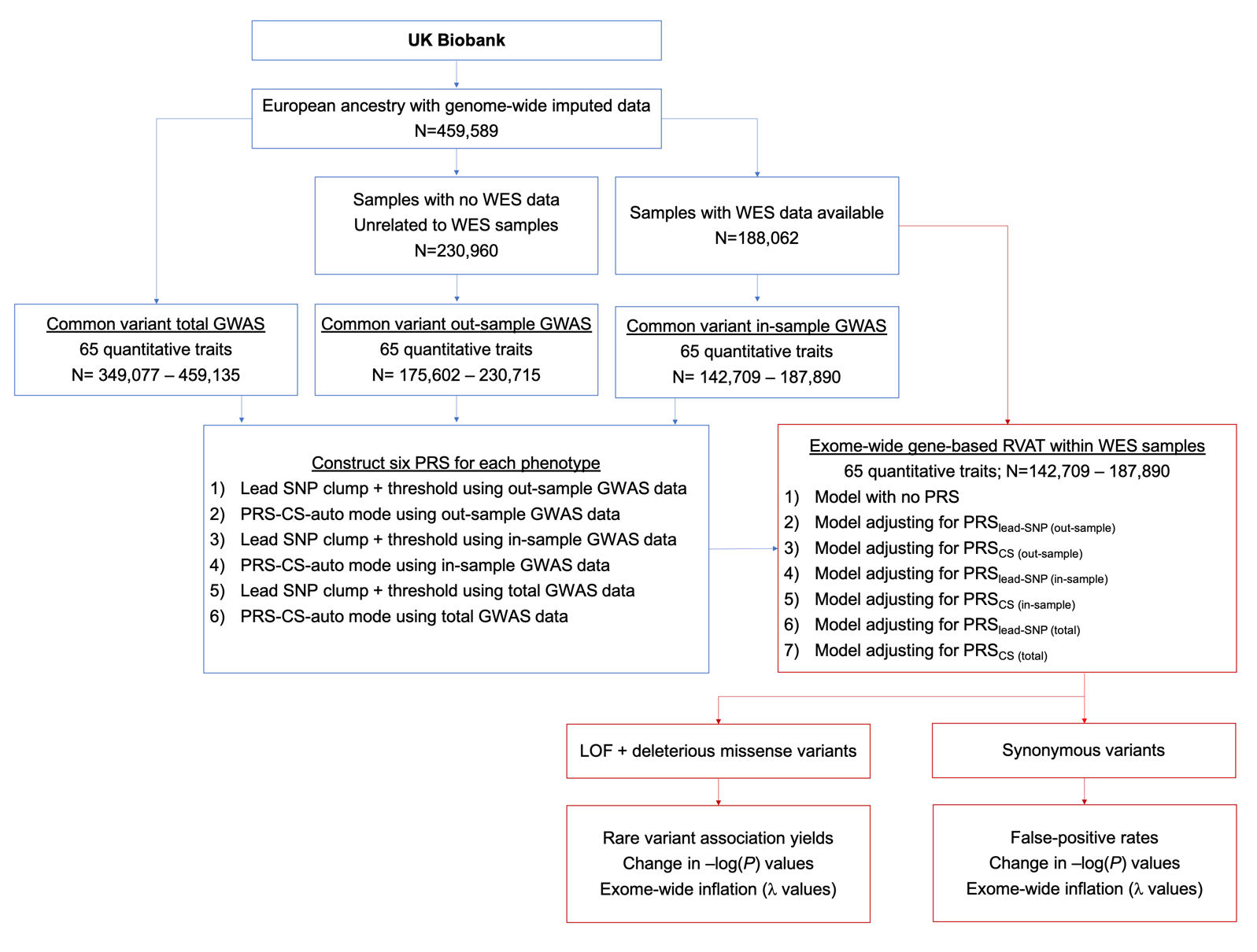

**Supplementary Figure 1: Analysis Flowchart.** The blue boxes show steps revolving around common variant analyses and PRS construction, while the red boxes highlight steps involving rare variant analyses.

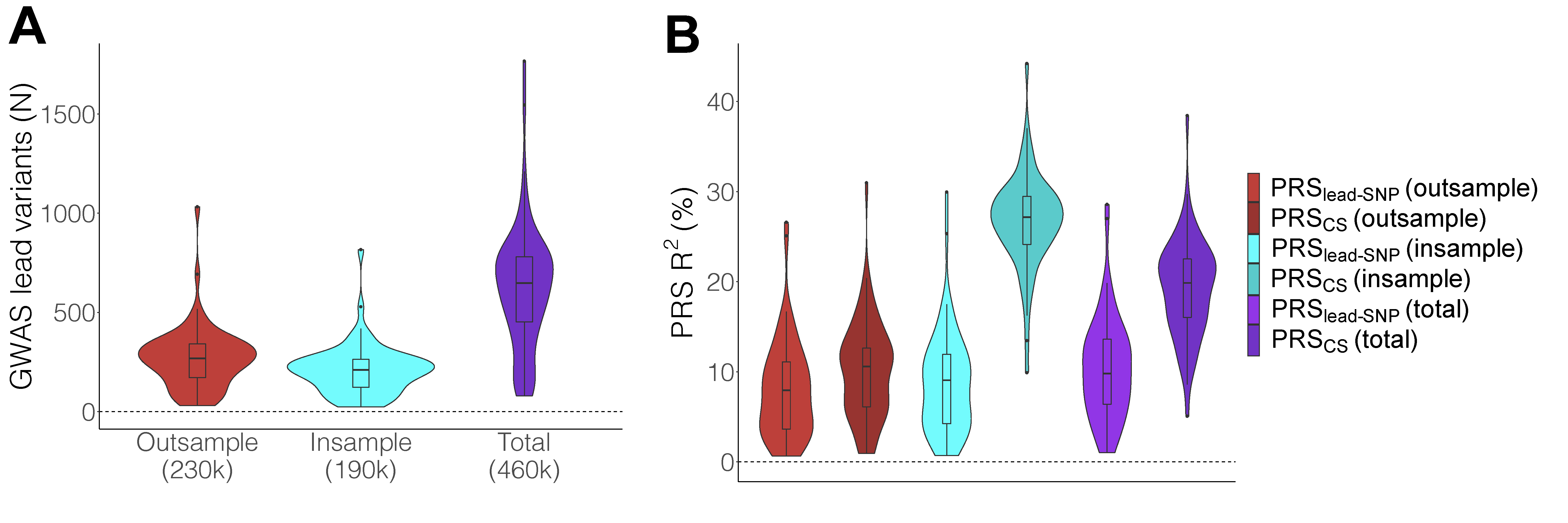

**Supplementary Figure 2: Number of significant lead variants from common variant GWAS and variance explained by subsequently derived PRS across the 65 traits. Panel A**: Violin plots for the number of significant independent lead variants from common variant GWAS across 65 phenotypes. Results from out-of-sample GWAS (230k, red), in-sample GWAS (190k, blue) and total GWAS (460k, purple) in the UK Biobank are shown. **Panel B**: Violin plots for the phenotypic variance explained by 6 types of PRS across the 65 phenotypes. Red shows two PRS derived from out-of-sample GWAS data, blue shows two PRS derived from completely in-sample GWAS data, while purple shows results for PRS derived from total GWAS data. All types of PRS explained variance for their respective traits, although we caution the interpretation of the magnitude of the R^2^ values for the in-sample and total PRS, as discovery samples were naturally also included in PRS testing. Boxplots: center line, median; box limits, upper and lower quartiles; whiskers, 1.5x interquartile range; points, outliers.

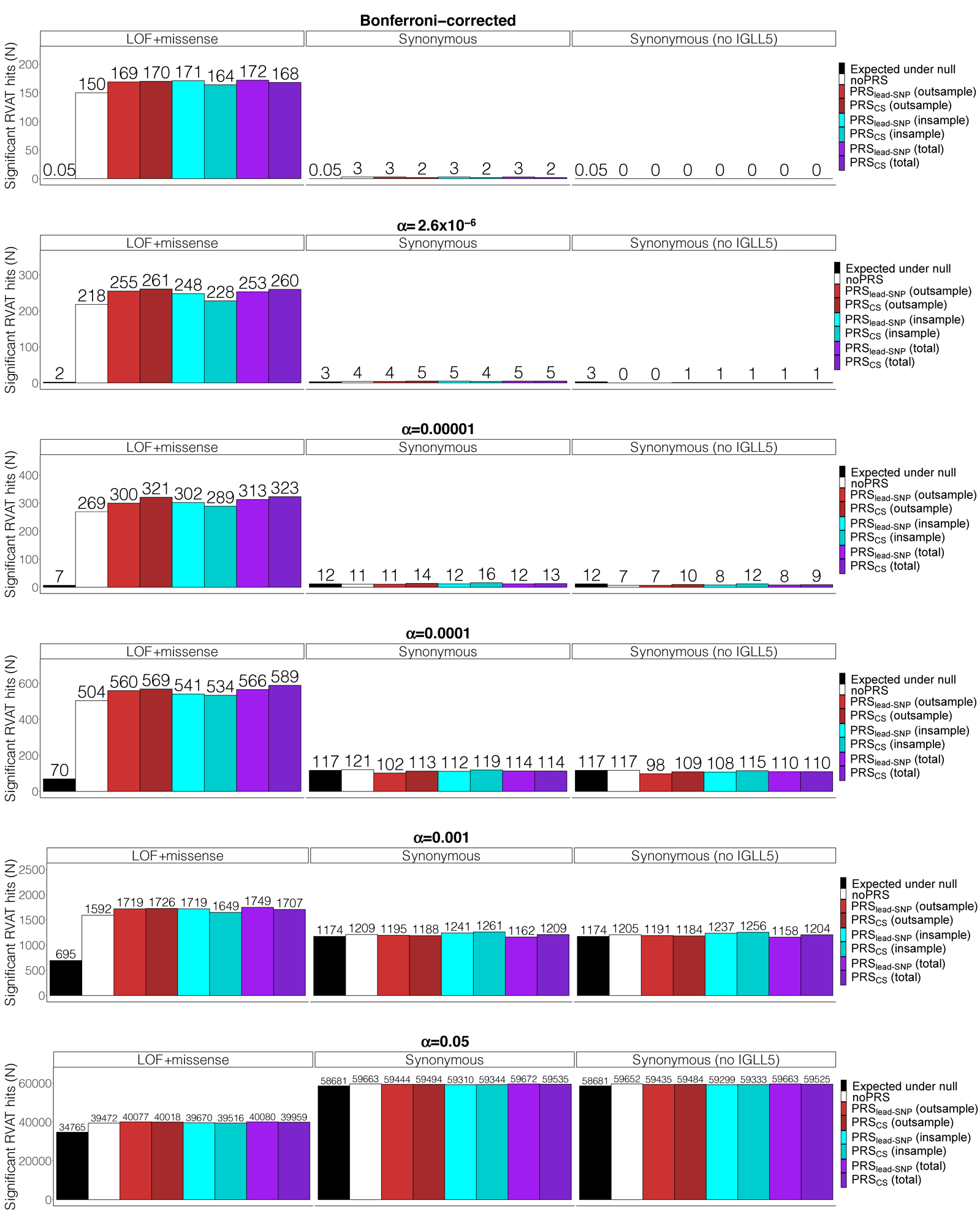

**Supplementary Figure 3: Bar plots for RVAT yield at different significance cutoffs.** Each row of plots shows results for a different significance threshold, with results from the analysis of deleterious variants (LOF and missense) plotted in the left panels and results for synonymous variants in the middle panels, and finally in the right panels the results for synonymous variants after excluding *IGLL5* gene-trait pairs. In each panel, the Y-axis shows the number of significant associations from RVAT. The black bars show the expected number of associations under the null, white bars show the number of associations using models without PRS, red bars show results adjusting for PRS based on out-sample GWAS, blue bars show results for PRS derived from in-sample GWAS, and purple shows results for total GWAS. Generally, PRS adjustment increases the number of identified associations strongly for the deleterious variant analysis - particularly at stringent cutoffs - while for synonymous variants the number of significant associations is similar between models without PRS and PRS adjusted models.

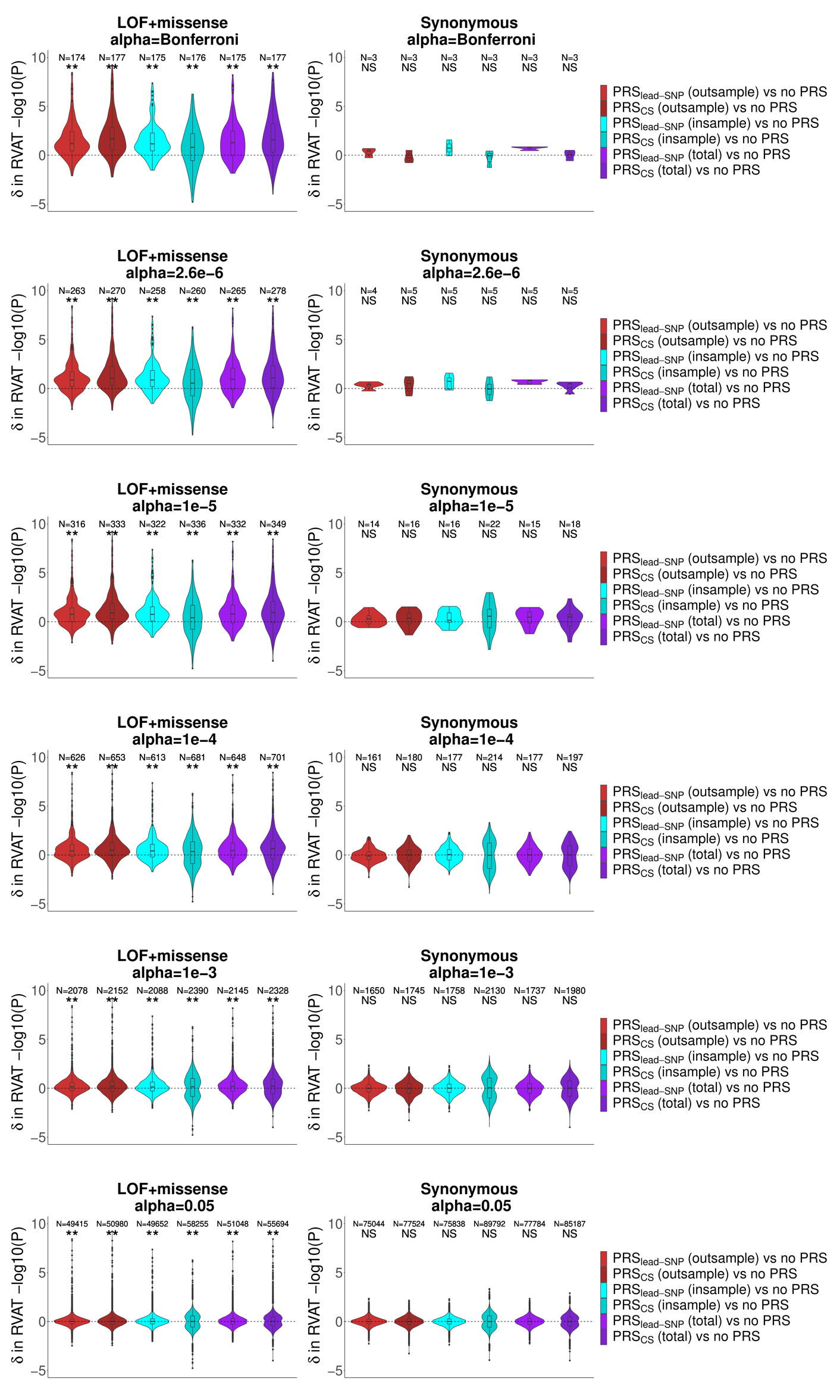

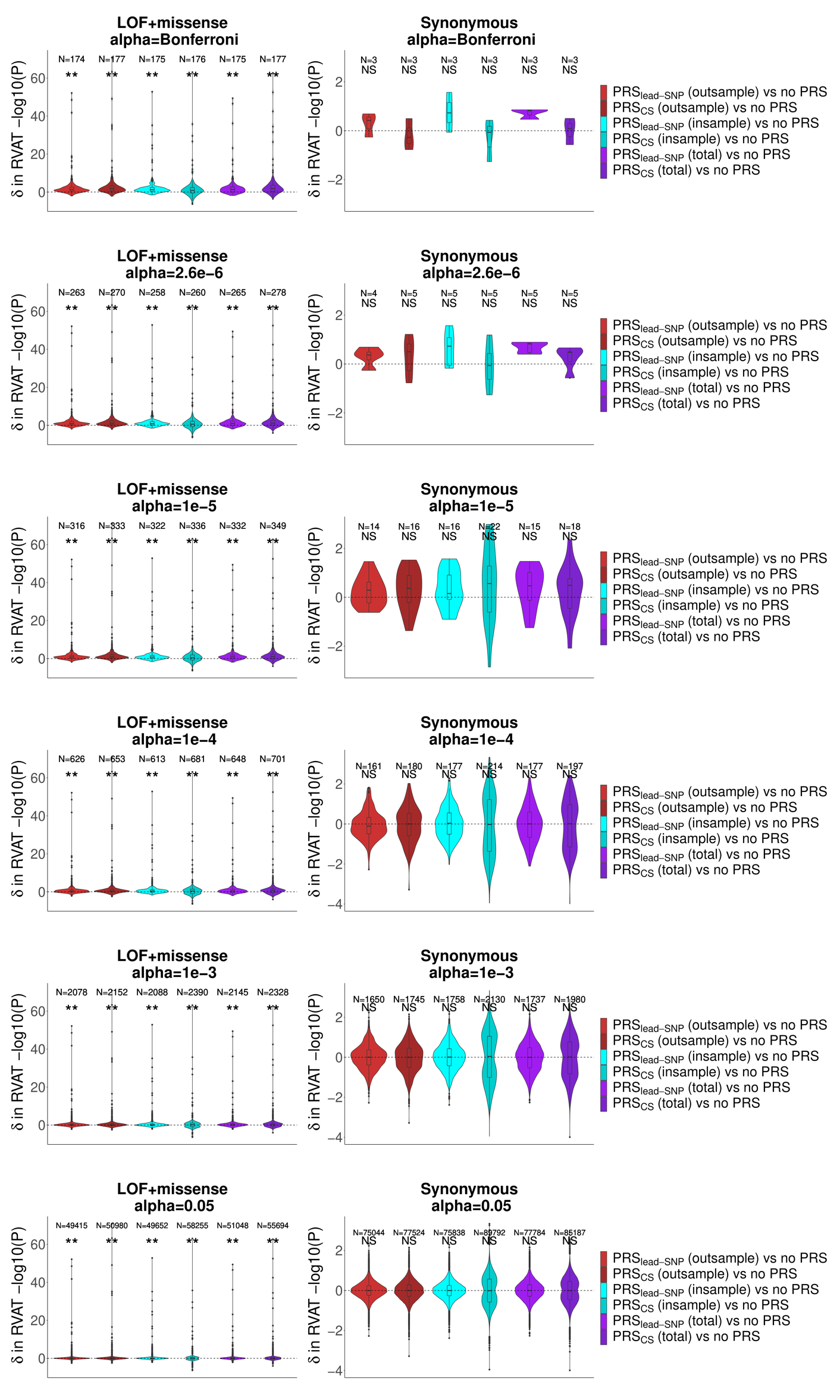

**A**

**B**

**Supplementary Figure 4: Violin plots for difference in *P*-values from models with PRS compared to models without PRS at different significance thresholds.** Each row of panels shows results analyzing *P*-values from gene-phenotype tests reaching significance at a different significance cutoff. Left panels always show results for deleterious variants (LOF and missense) while right panels show results for synonymous variants. The difference (δ) in -log10(P) values for gene-phenotype tests between a model including PRS as compared to the model without PRS are plotted on the y-axis. Red shows results from PRS based on out-of-sample GWAS data, blue shows results for PRS derived from in-sample GWAS data, and purple shows results for PRS derived from total GWAS data. ** indicates strongly significant (positive d̅ and *P*<0.0014 = 0.05 / [6 thresholds*6 models]) using two-sided paired Wilcoxon-rank tests; * indicates suggestively significant (positive d̅ and *P*<0.0083 = 0.05 / 6 thresholds); NS indicates not significant (negative d̅ or *P*>0.0083). **Part A** and **Part B** show the same data, where **Part A** is equally scaled across panels for clarity (and therefore outlier data points are not plotted), while **Part B** shows all data points and therefore y-axes are not equally scaled across panels. Boxplots: center line, median; box limits, upper and lower quartiles; whiskers, 1.5x interquartile range; points, outliers.

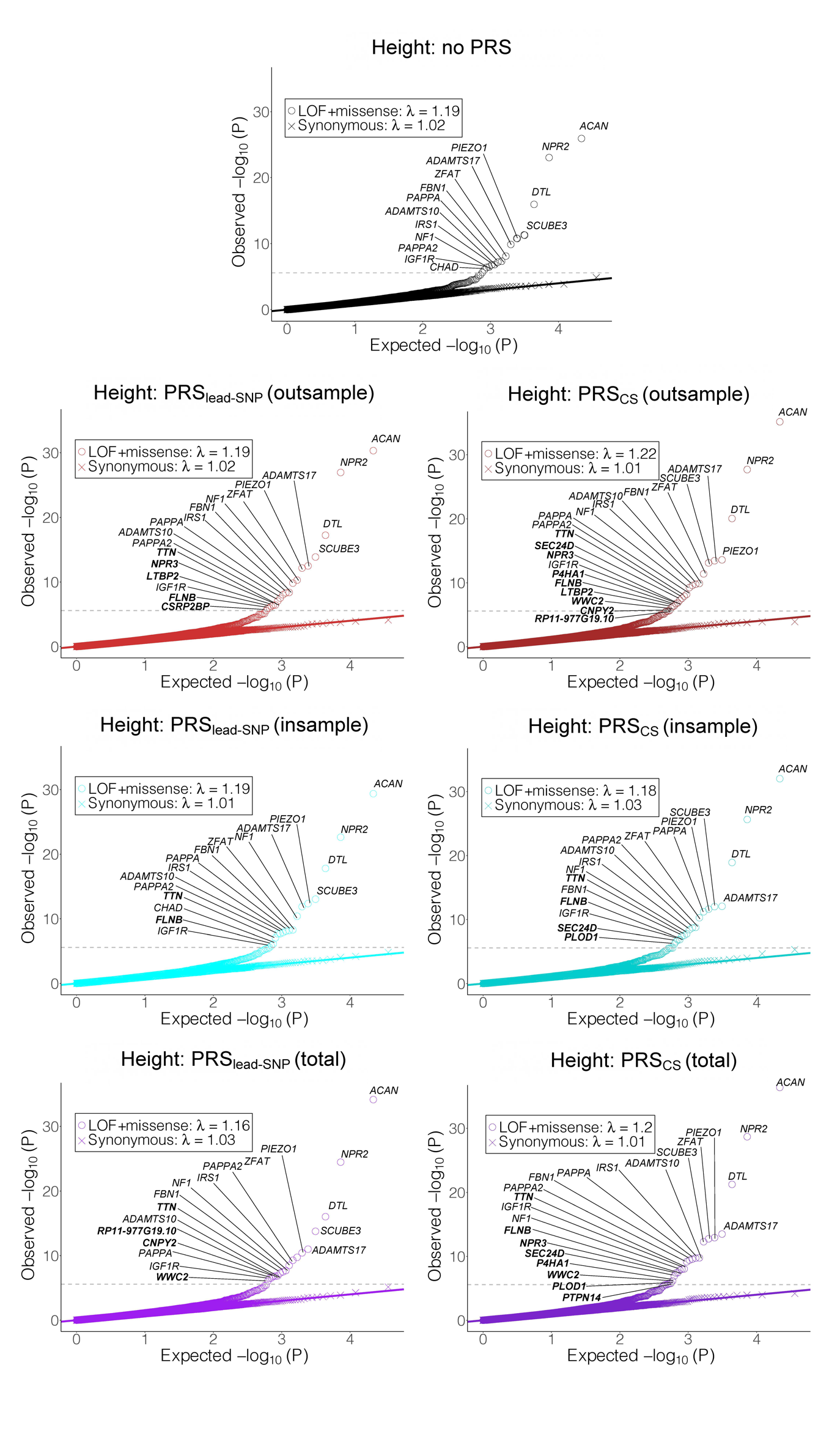

**Supplementary Figure 5: Quantile-quantile plots for the phenotype height with exome-wide significant genes annotated.** In each plot, the y-axis shows the *P*-values from exome-wide RVAT, while the x-axis shows the *P*-values expected under the null. QQ-plots for deleterious variants (LOF+missense) are plotted using circles, while QQ-plots synonymous variants are plotted using crosses. Black points show the results for the model without PRS, red shows the results for the model adjusted for PRS derived out-of-sample GWAS data, blue shows results for PRS derived from in-sample GWAS data, and purple shows results for PRS derived from total GWAS data. Genes reaching exome-wide significance (*P*<2.6x10^-6^) are annotated with the gene name; genes that only reached significance after PRS adjustment are shown in bold. In all cases, more genes were identified after PRS adjustment for this phenotype; only one gene, *CHAD*, dropped below exome-wide significance after PRS adjustment for certain models.

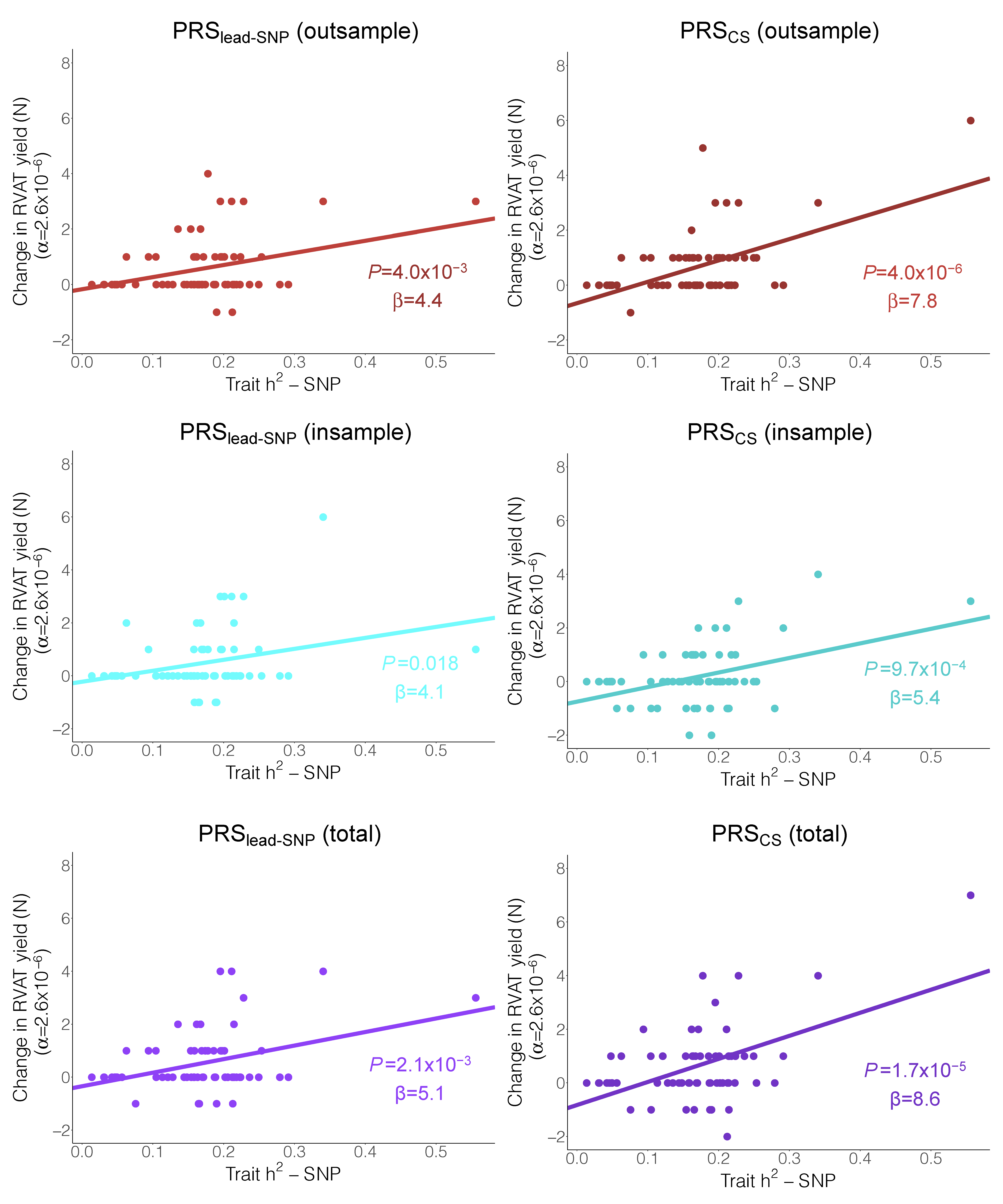

**Supplementary Figure 6: Correlation between SNP-heritability and the change in the number of significant rare variant associations after PRS adjustment across the 65 traits.** In each plot, the x-axis represents trait SNP-heritability (h^2^_SNP_) estimated using Linkage Disequilibrium Score Regression. The y-axis represents the change in the number of RVAT associations reaching exome-wide significance (α=2.6x10^-6^) after adjusting for PRS, across the studied traits (N=65). RVAT yield change (defined as the difference in the number of significant associations after PRS adjustment compared to models without PRS) is regressed on h^2^_SNP_ using ordinary linear regression; the regression trend line is added in each plot. For all models, there is a trend towards a positive association between trait h^2^_SNP_ and change in RVAT yield (*P*<0.05 and β>0). The trend reached *P<*0.0083 (=0.05/6) for all PRS models except PRS_CS (insample)_.

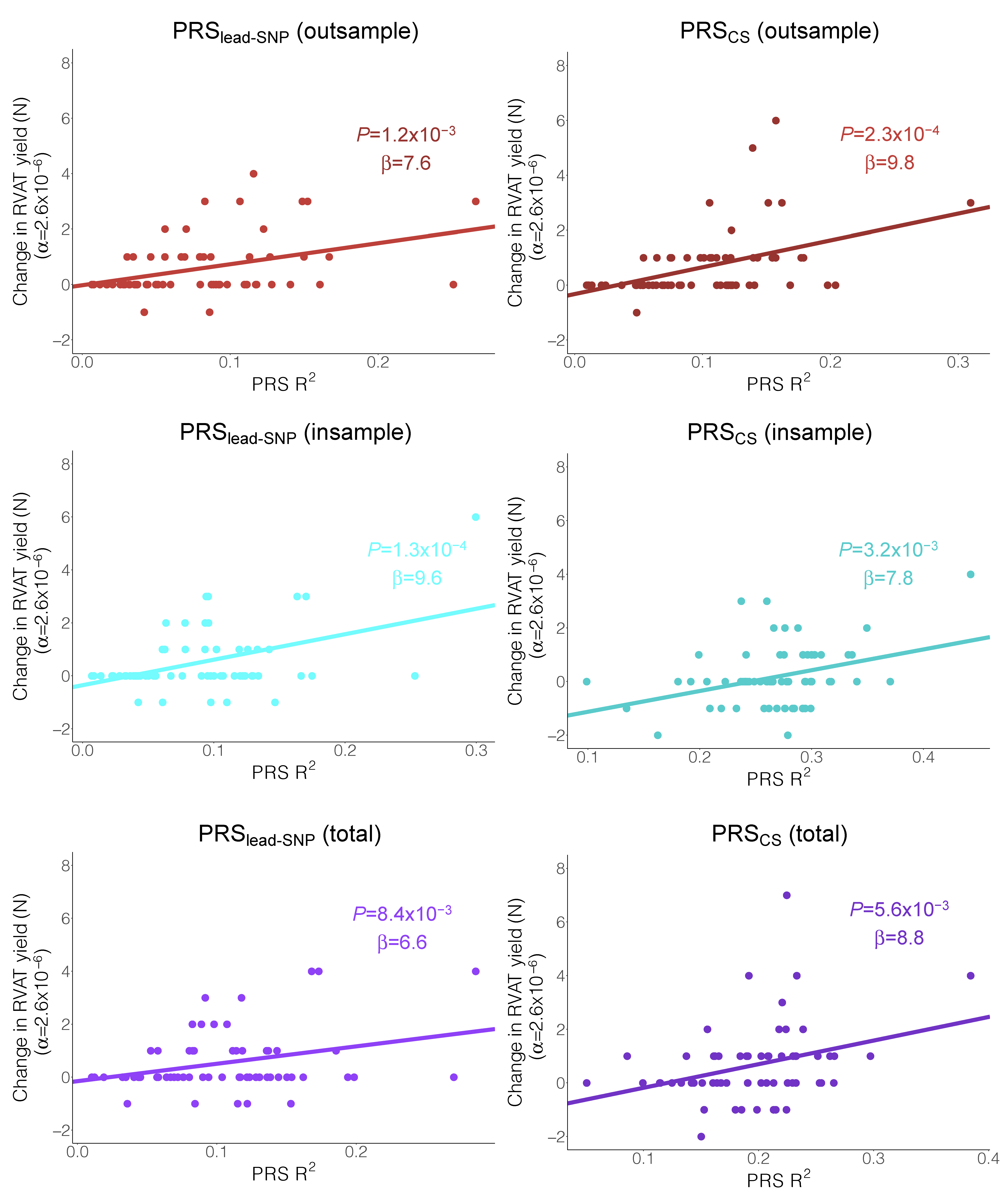

**Supplementary Figure 7: Correlation between PRS variance explained and the change in the number of significant rare variant associations after PRS adjustment across the 65 traits.** In each plot, the x-axis represents the variance explained (R^2^) of the PRS for its respective trait in the exome sequencing cohort. The y-axis represents the change in the number of RVAT associations reaching exome-wide significance (α=2.6x10^-6^) after adjusting for PRS, across the studied traits (N=65). RVAT yield change (defined as the difference in the number of significant associations after PRS adjustment compared to models without PRS) is regressed on the R^2^ using ordinary linear regression; the regression trend line is added in each plot. For all models, there is a trend towards a positive association between PRS R^2^ and change in RVAT yield (*P*<0.05 and β>0). The trend reached *P<*0.0083 (=0.05/6) for all PRS models except PRS_lead-SNP (total)_ (*P*=0.0084).

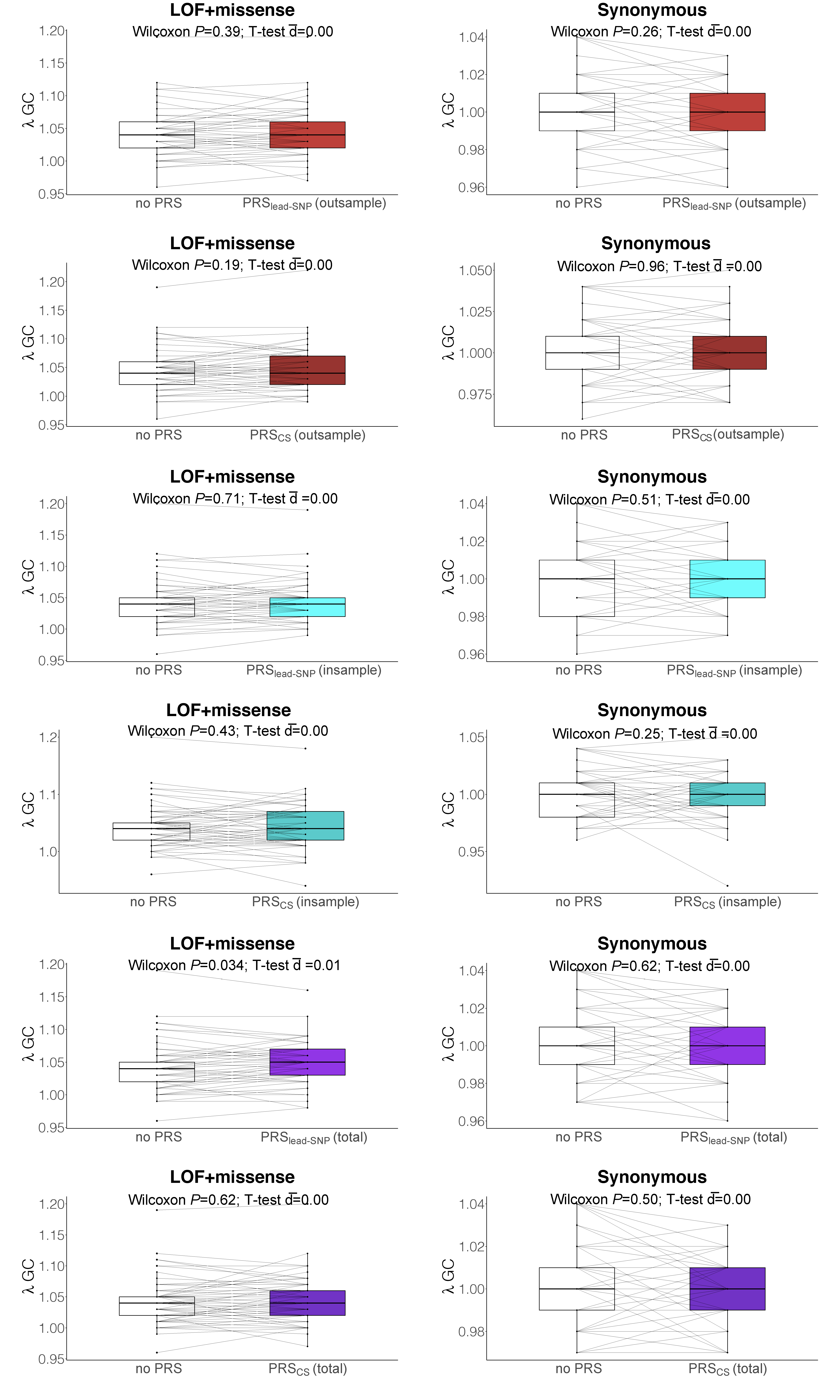

**Supplementary Figure 8: Paired box plots for comparison of genomic inflation factors between models without PRS and models with PRS.** Left panels show results for deleterious variants (LOF and missense) while right panels show results for synonymous variants. Y-axes show the genomic inflation factors, λ_GC_, from exome-wide analyses. In each plot, the white box plot shows genomic inflation factors from models without PRS, while red show results from PRS derived from out-of-sample GWAS data, blue shows results for PRS derived from in-sample GWAS, and purple shows results for PRS derived from total GWAS. Comparisons are annotated with *P*-values from two-sided paired Wilcoxon signed rank tests, as well estimated differences (d̅) in λ_GC_ between models with PRS and models without PRS estimated from paired T-tests after removing outliers. Boxplots: center line, median; box limits, upper and lower quartiles; whiskers, 1.5x interquartile range; points, outliers.

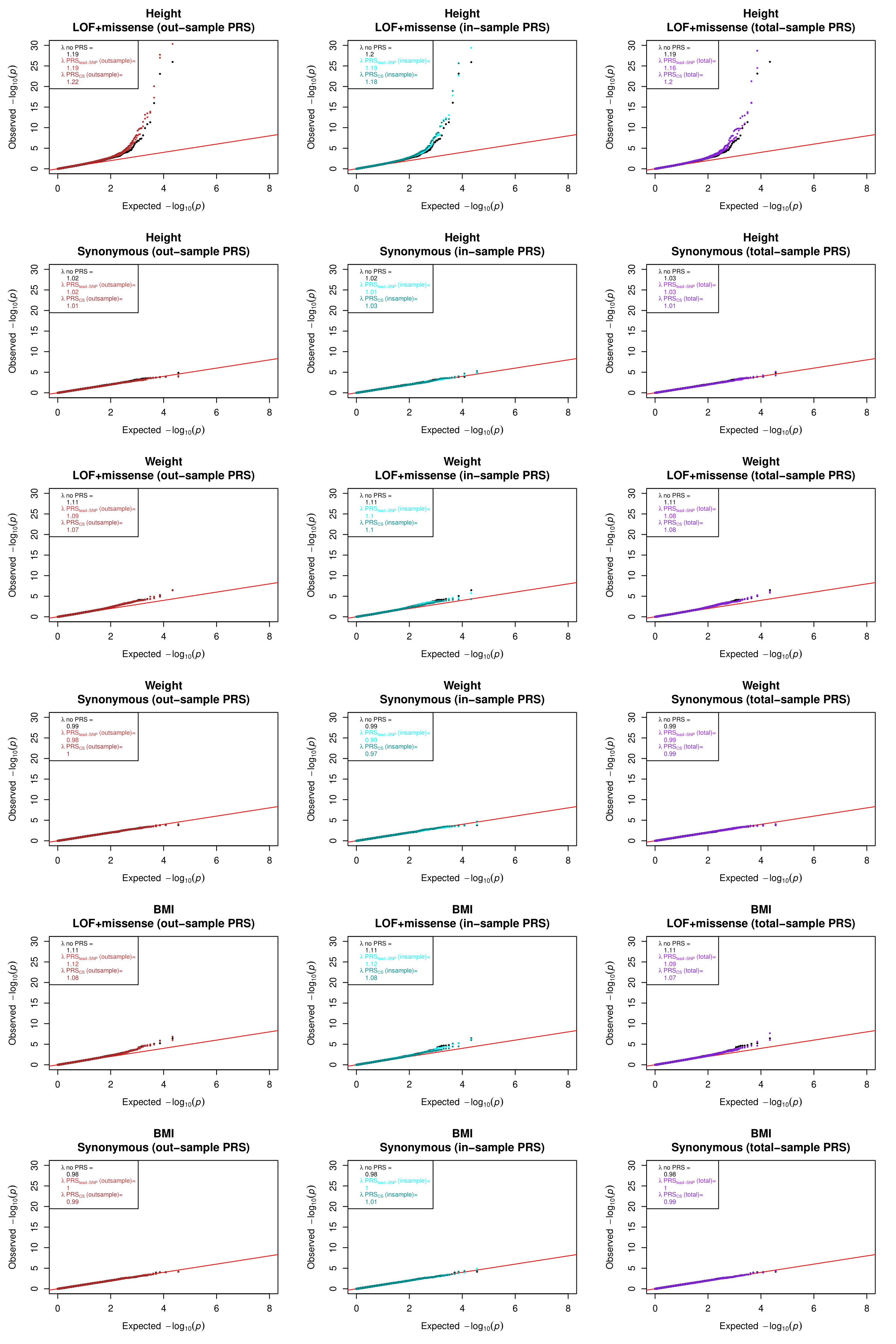

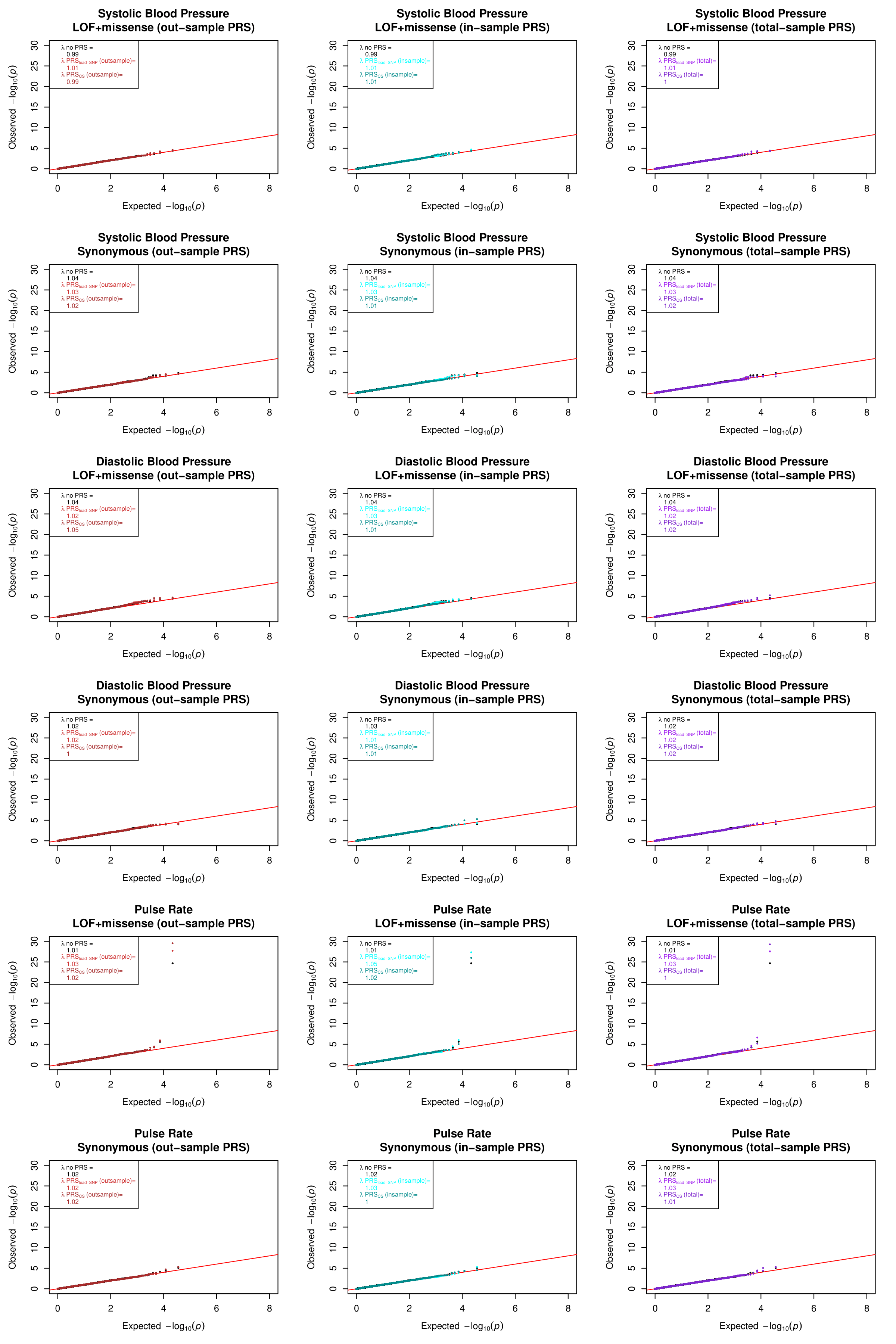

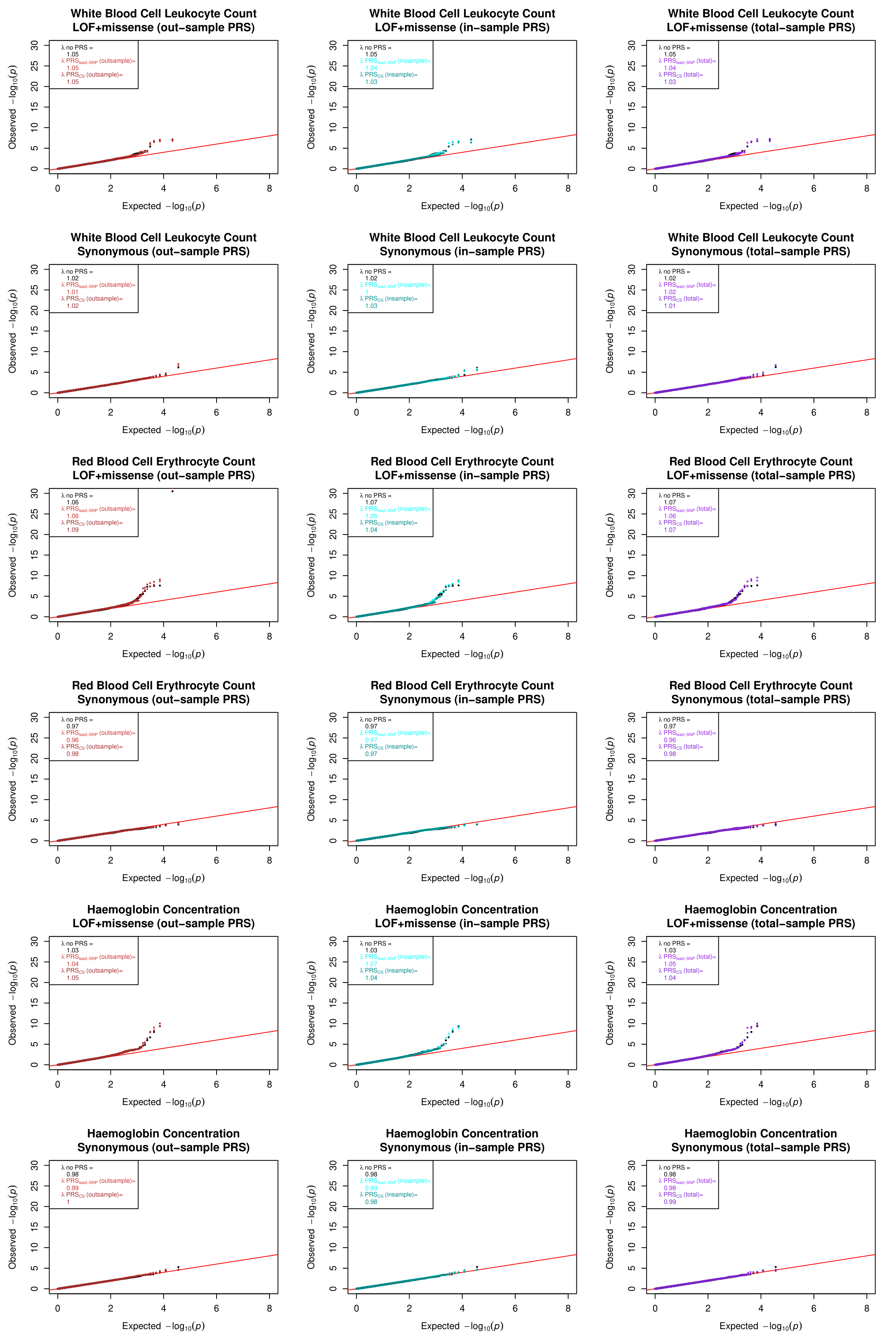

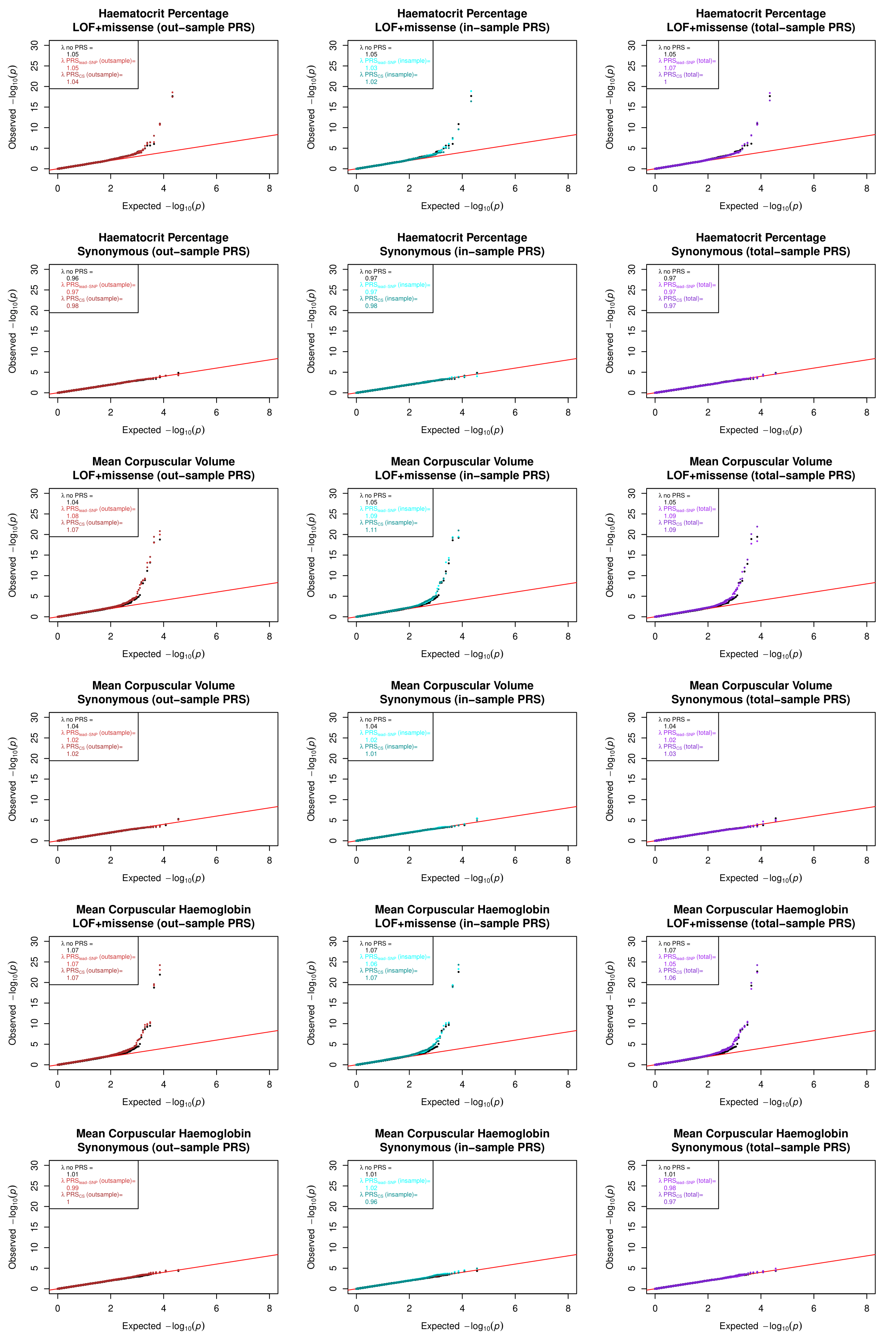

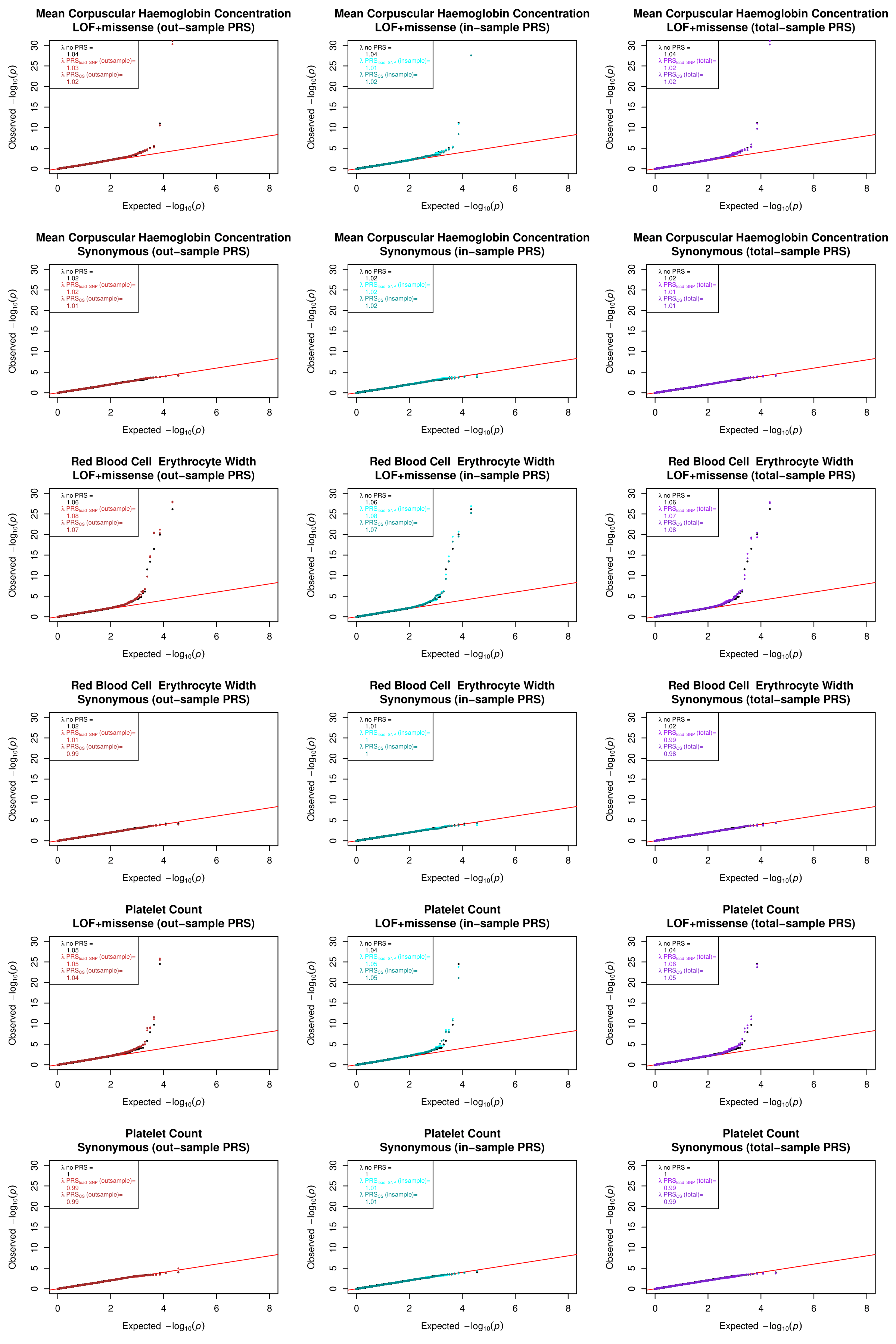

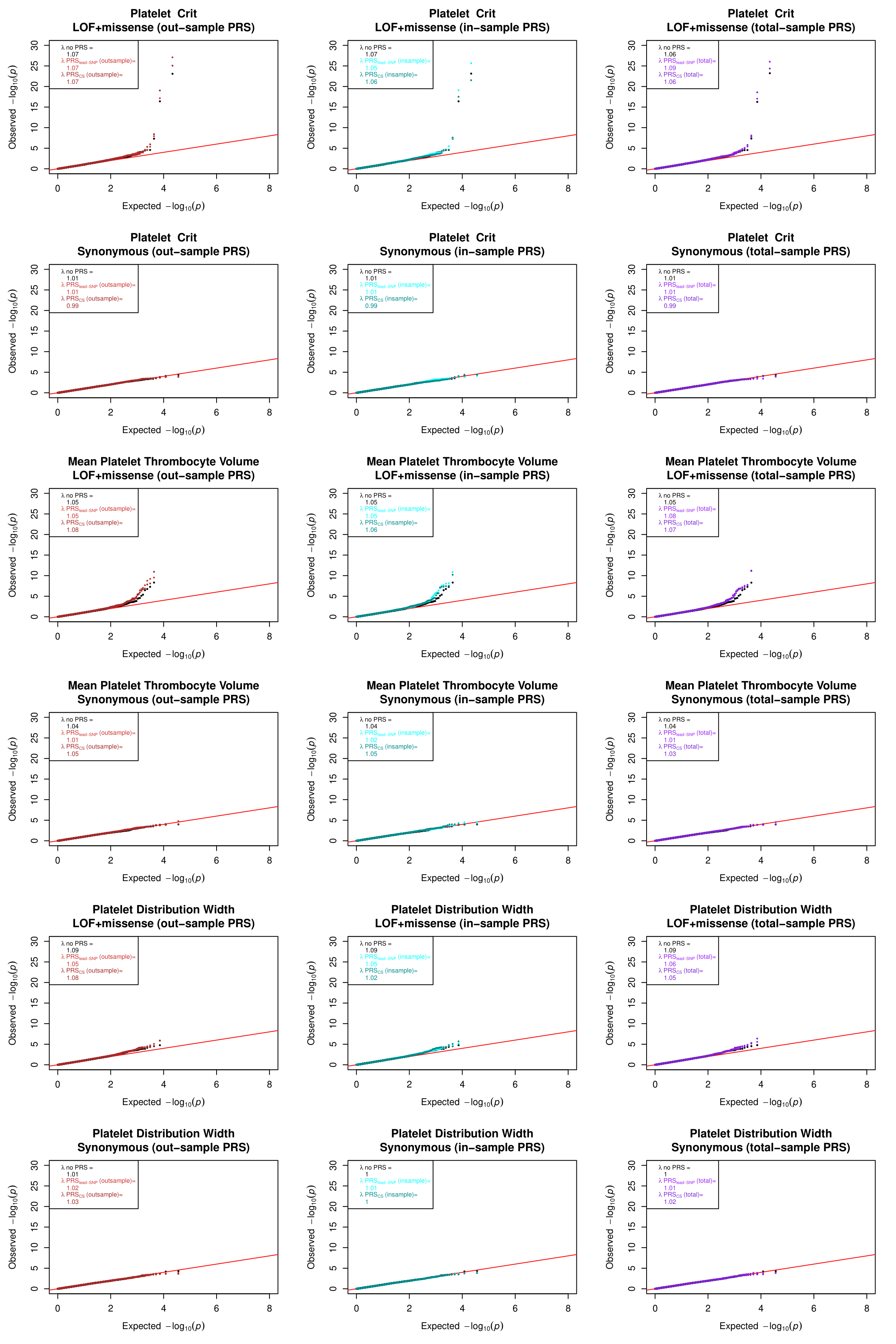

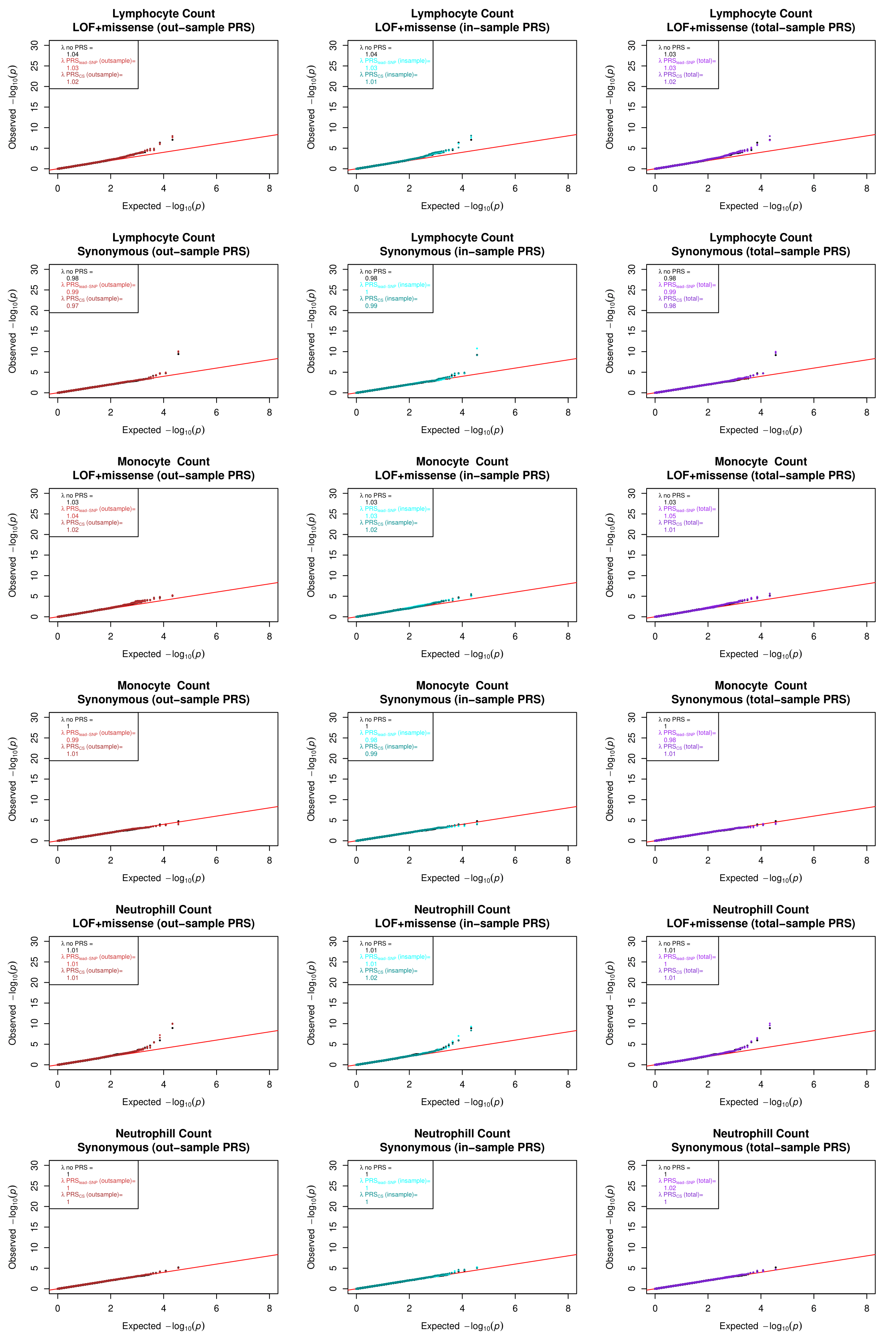

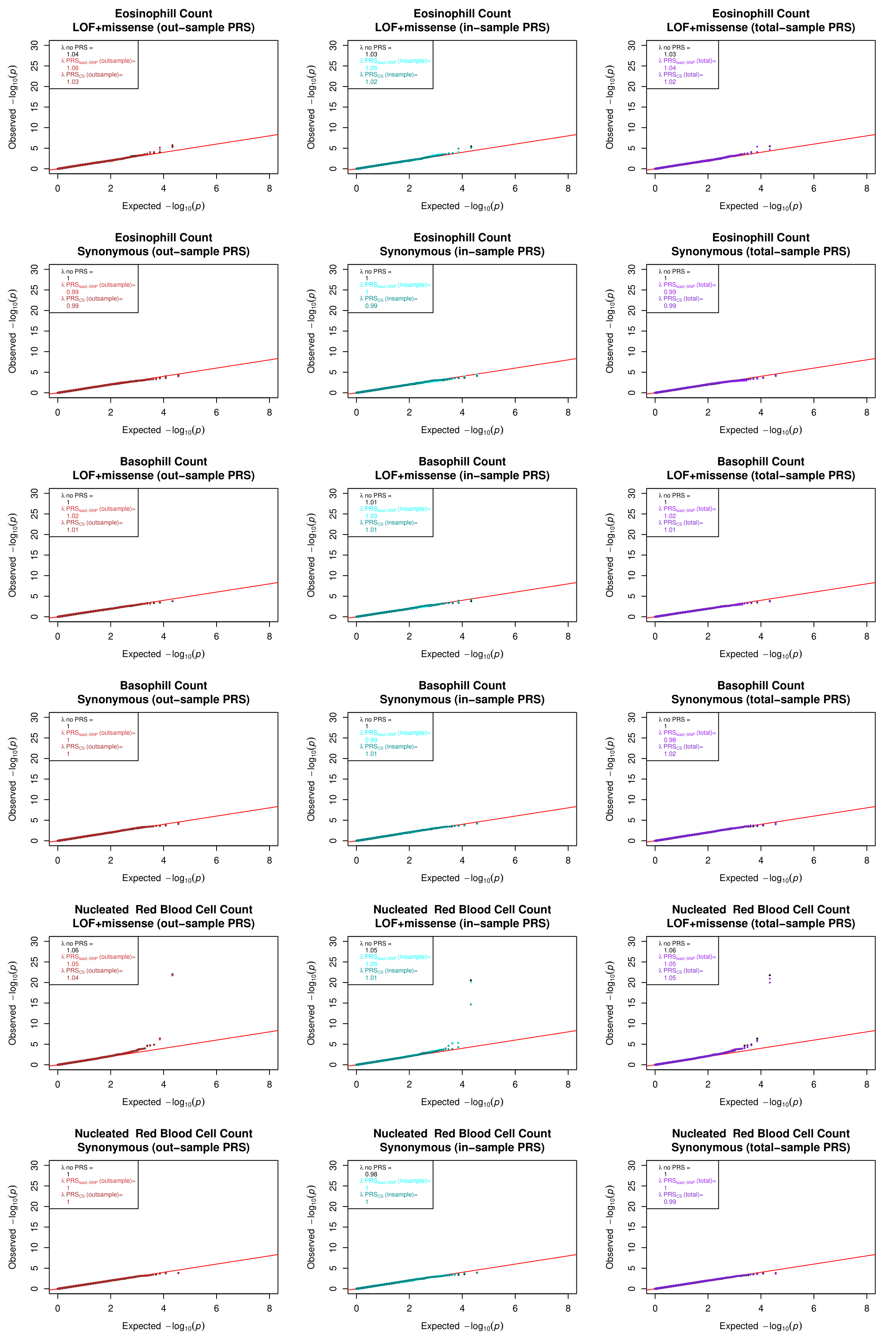

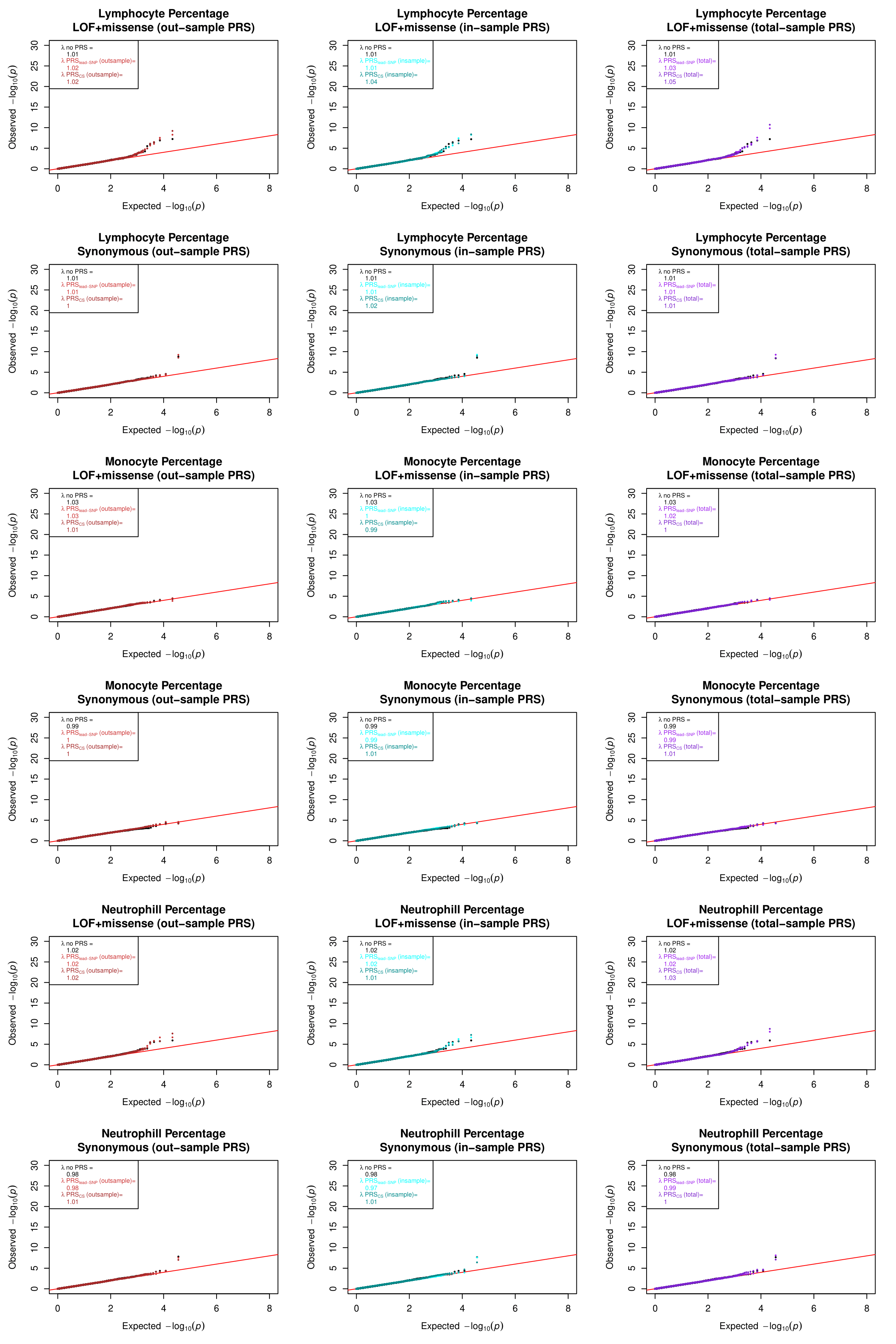

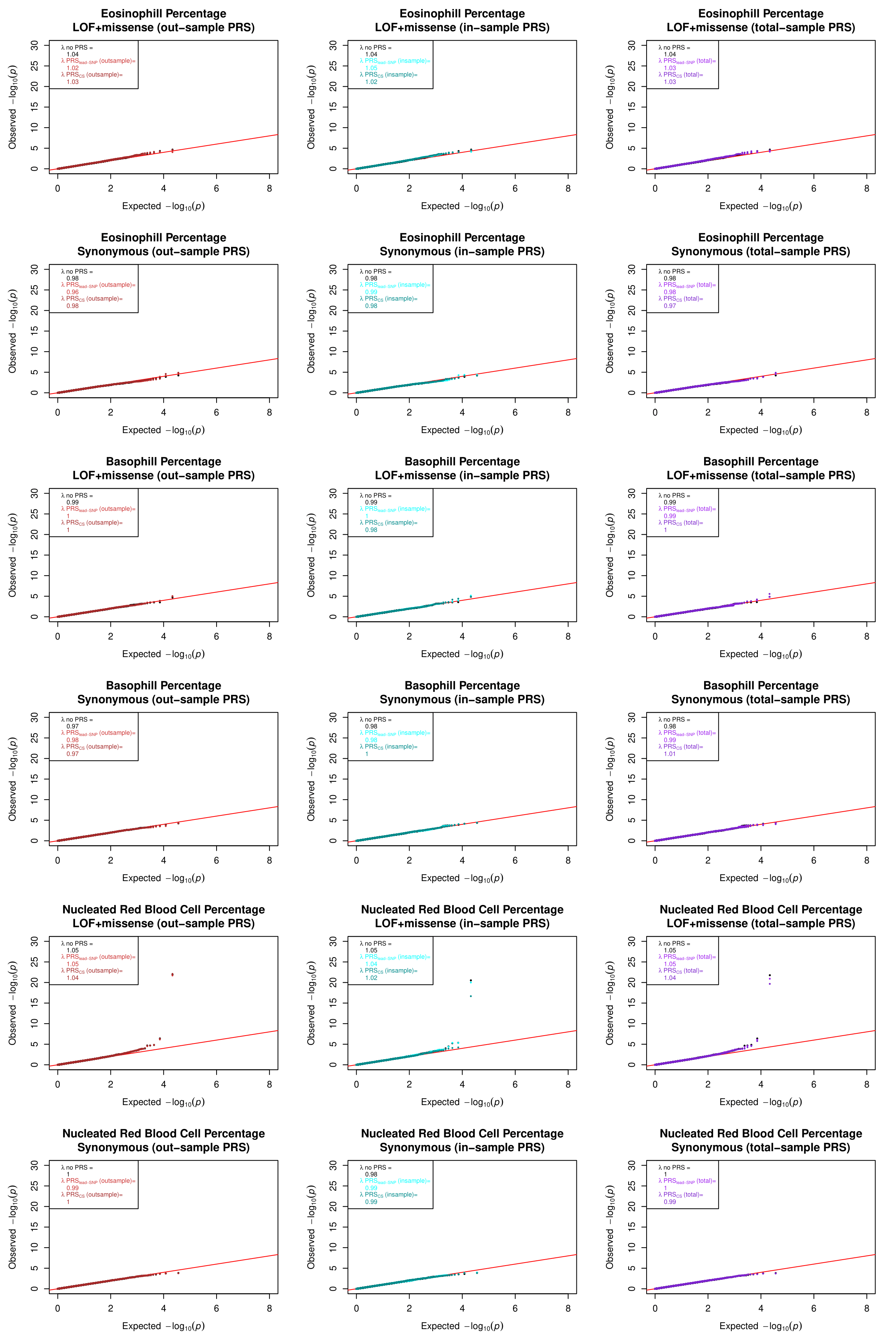

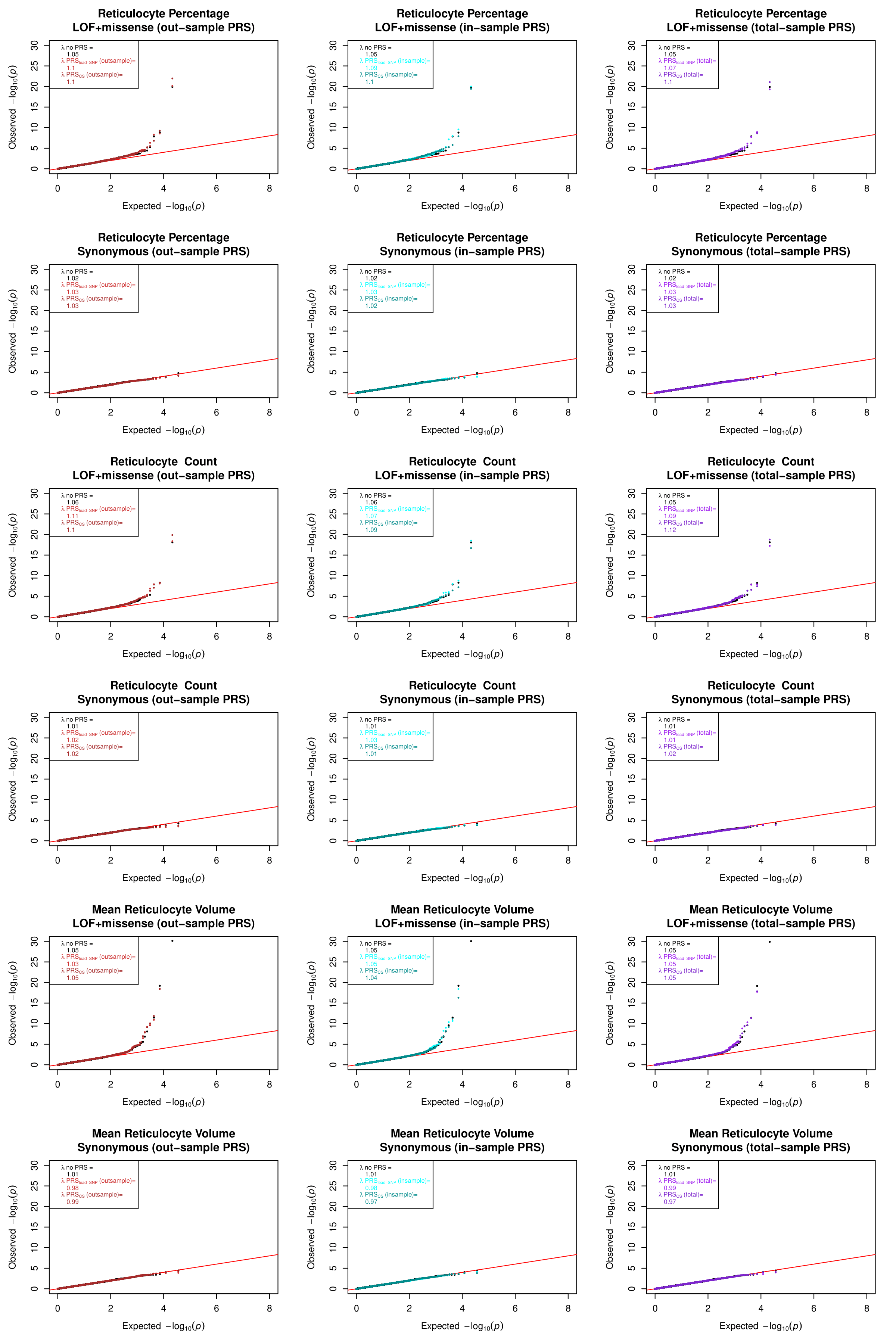

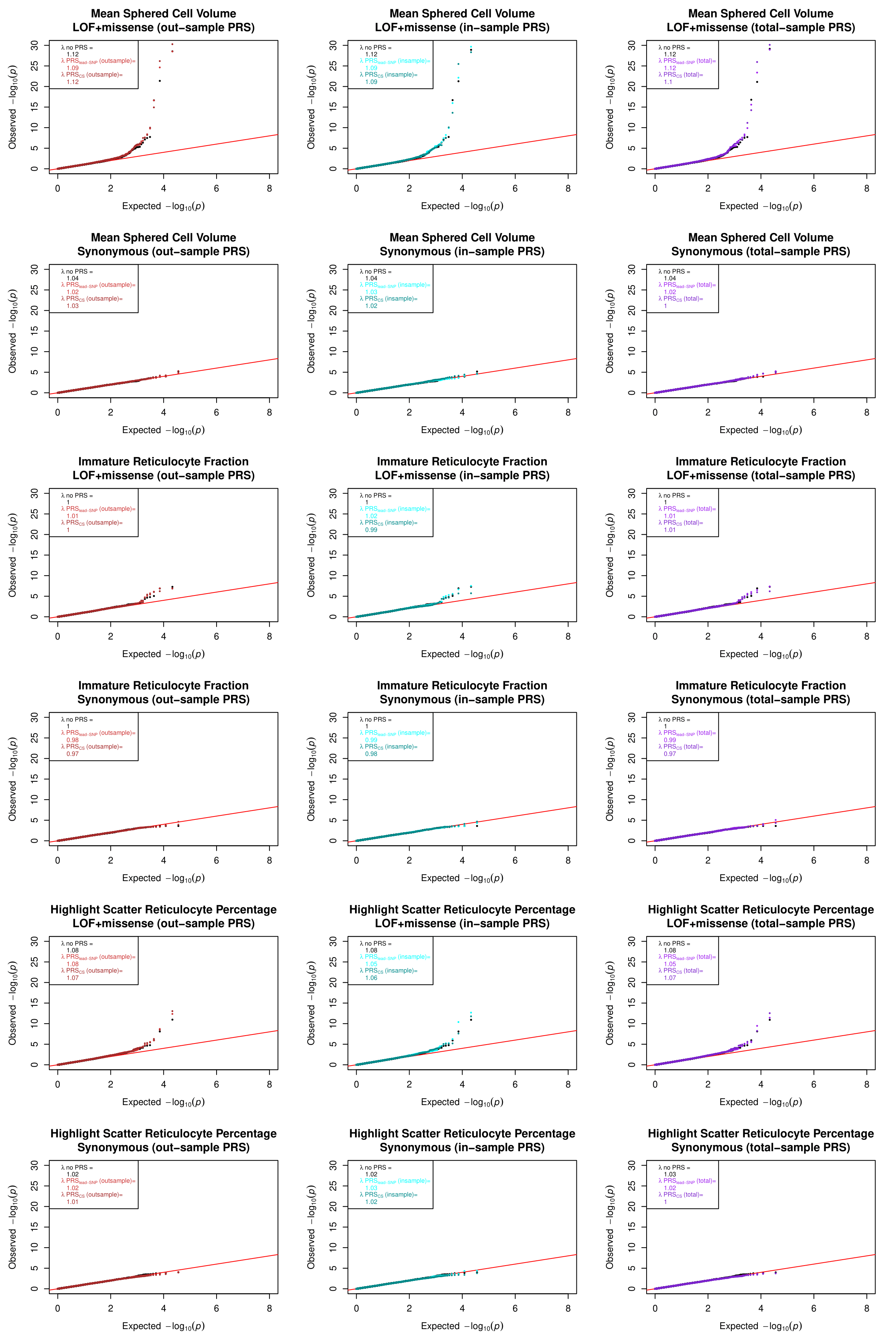

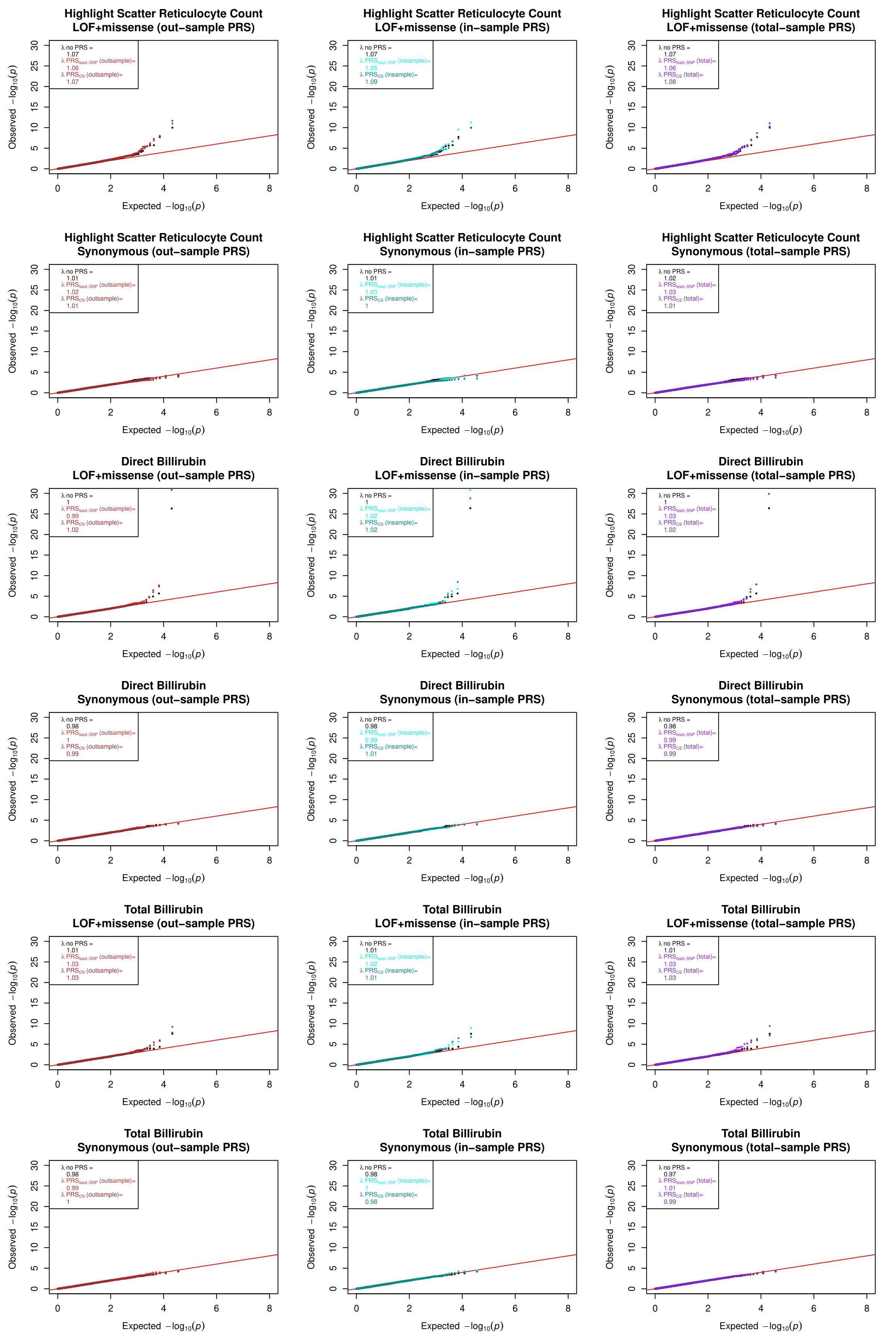

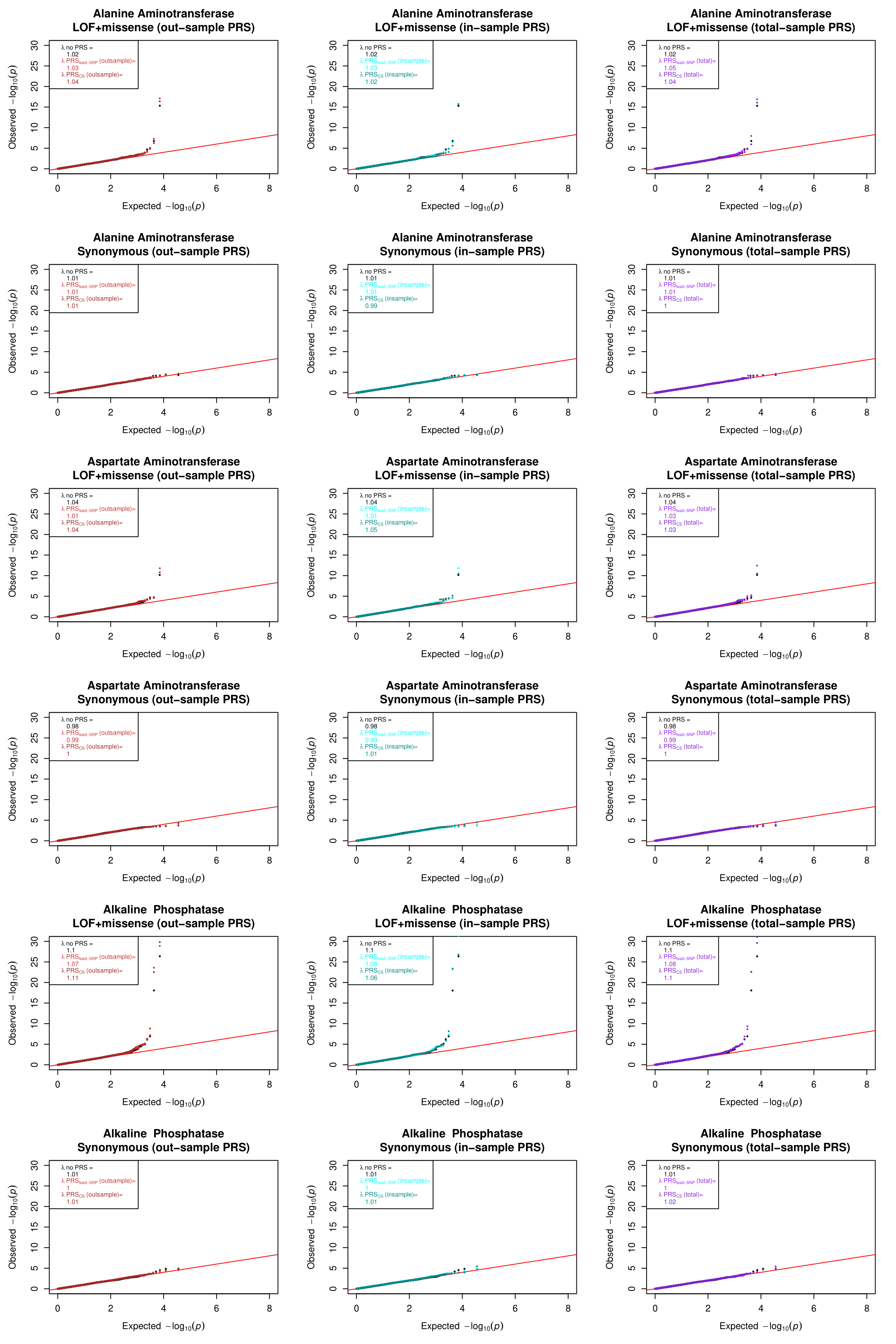

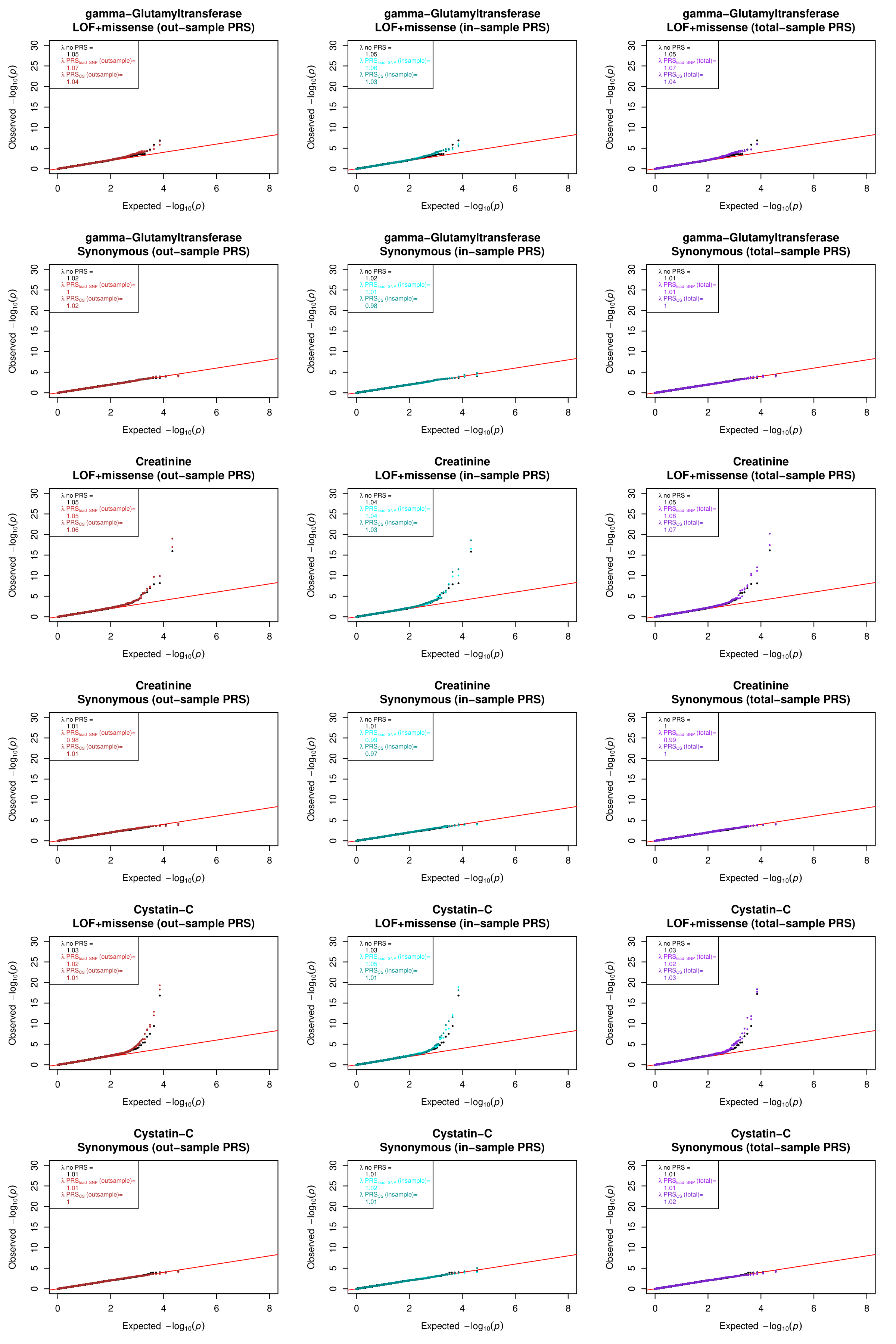

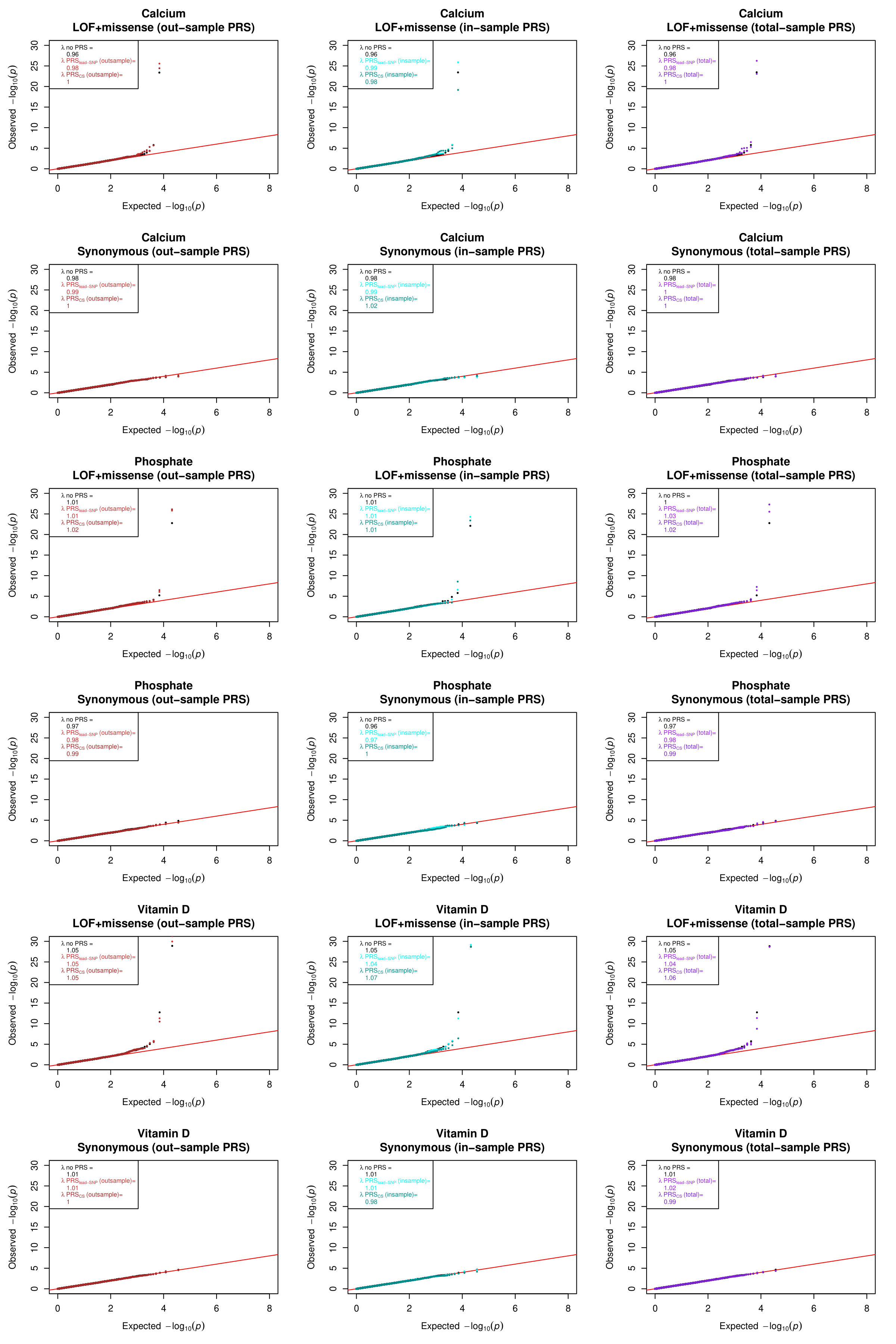

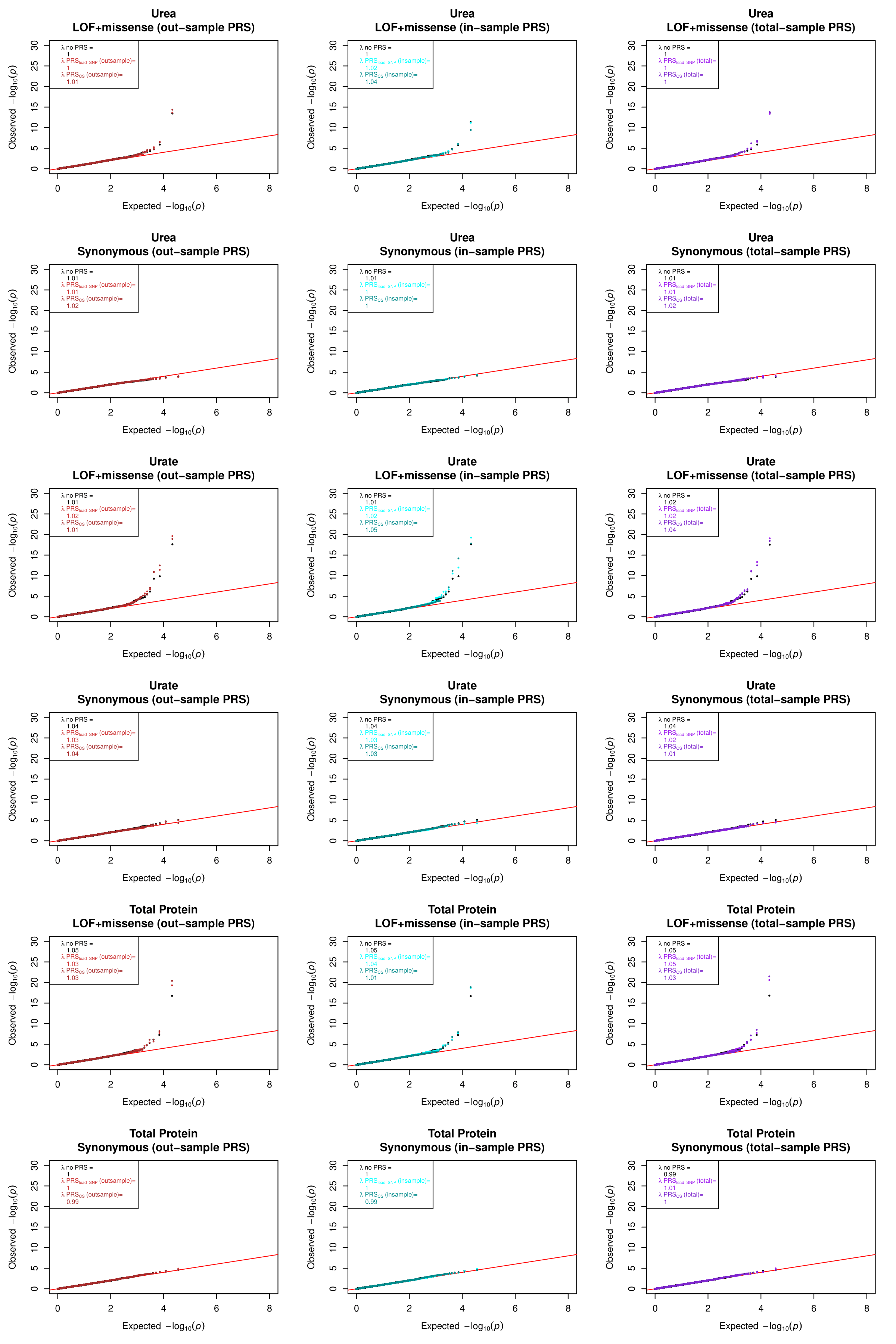

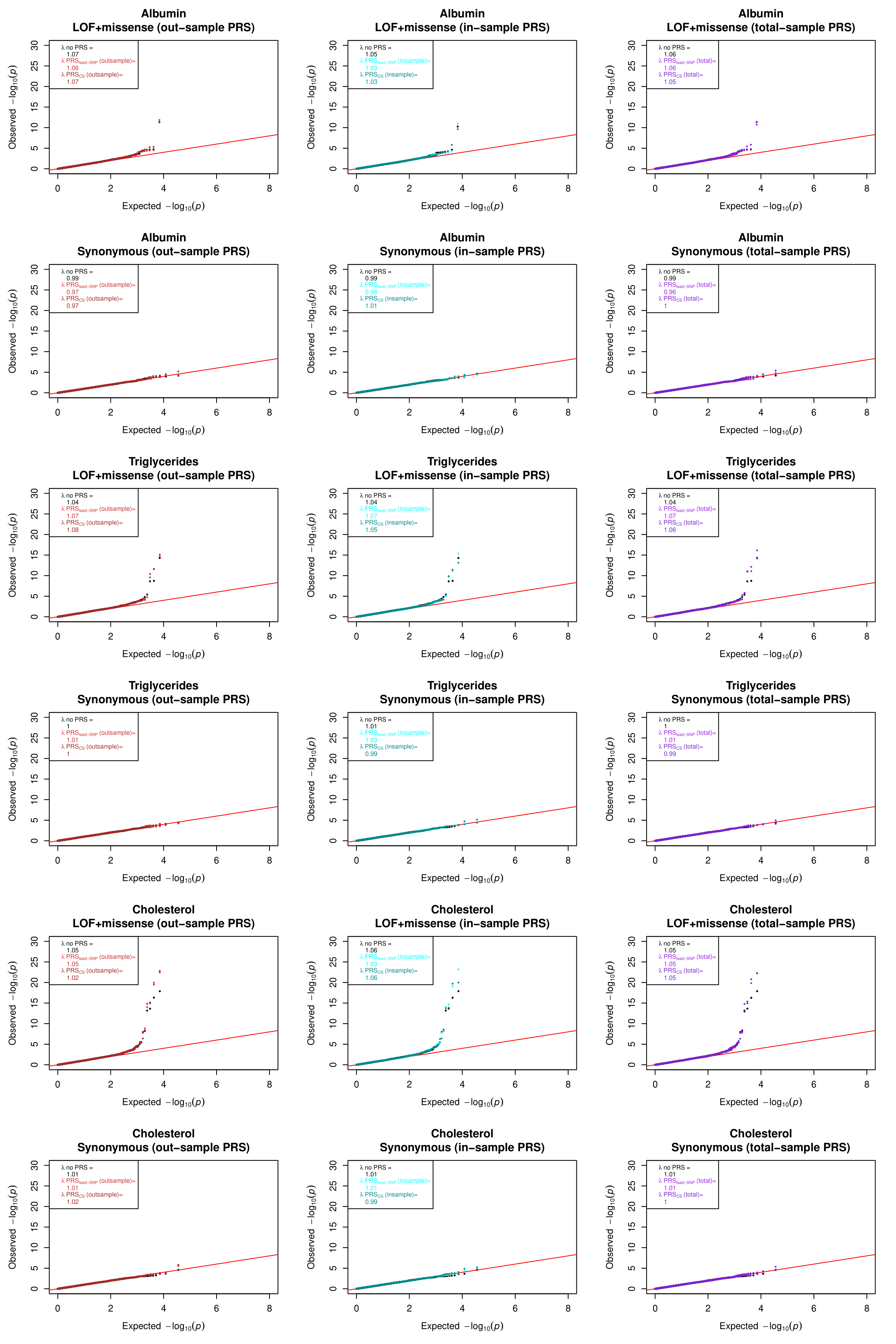

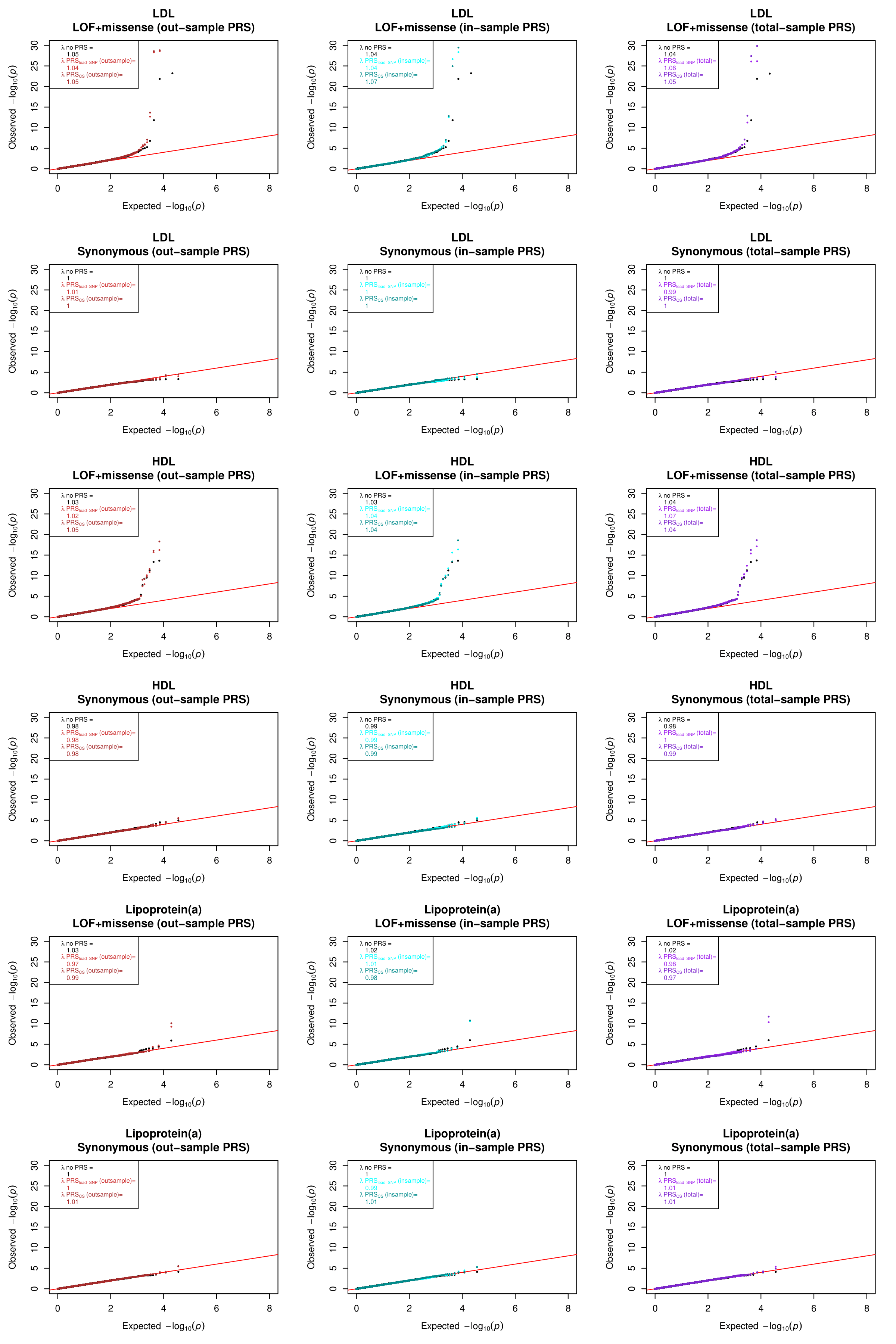

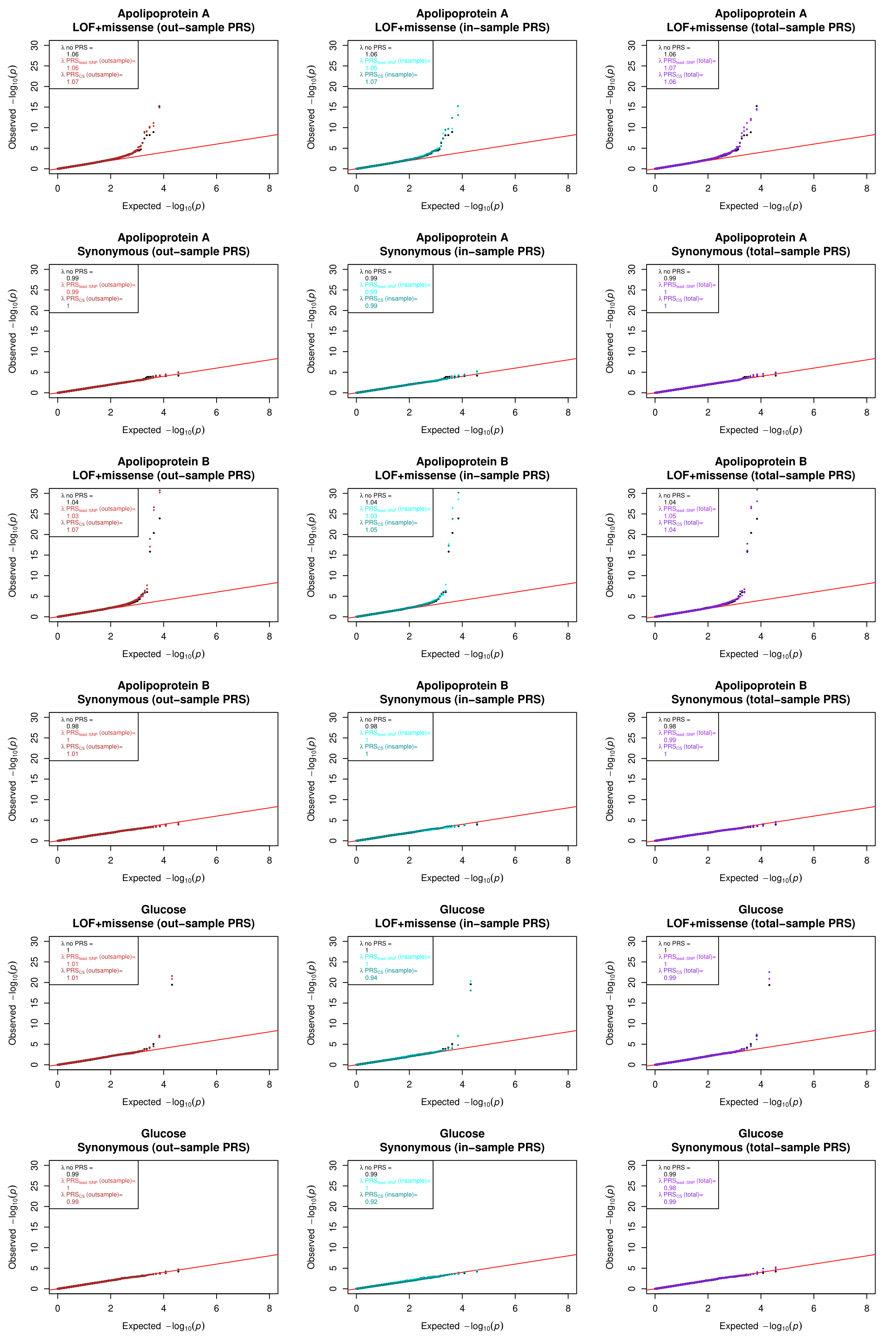

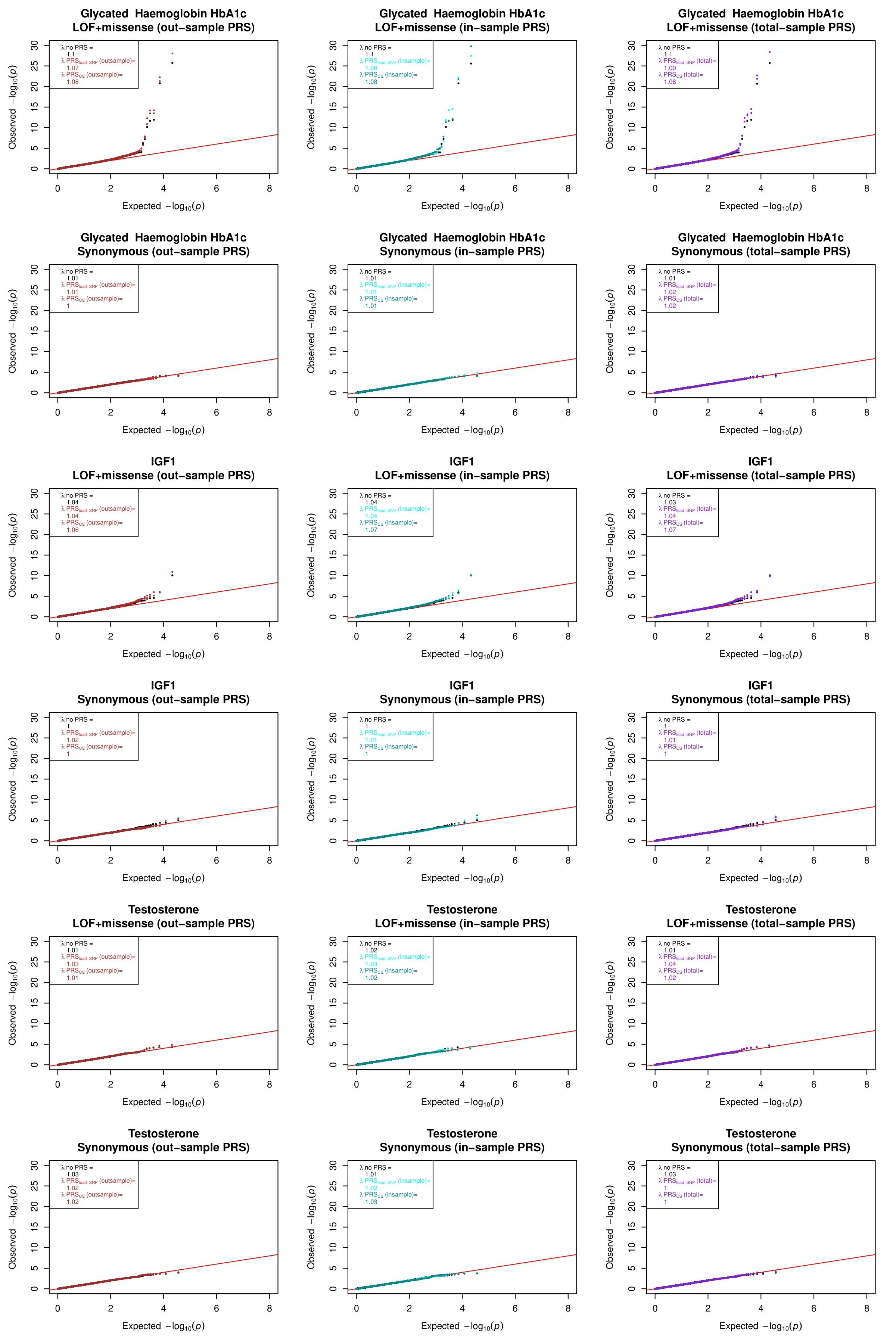

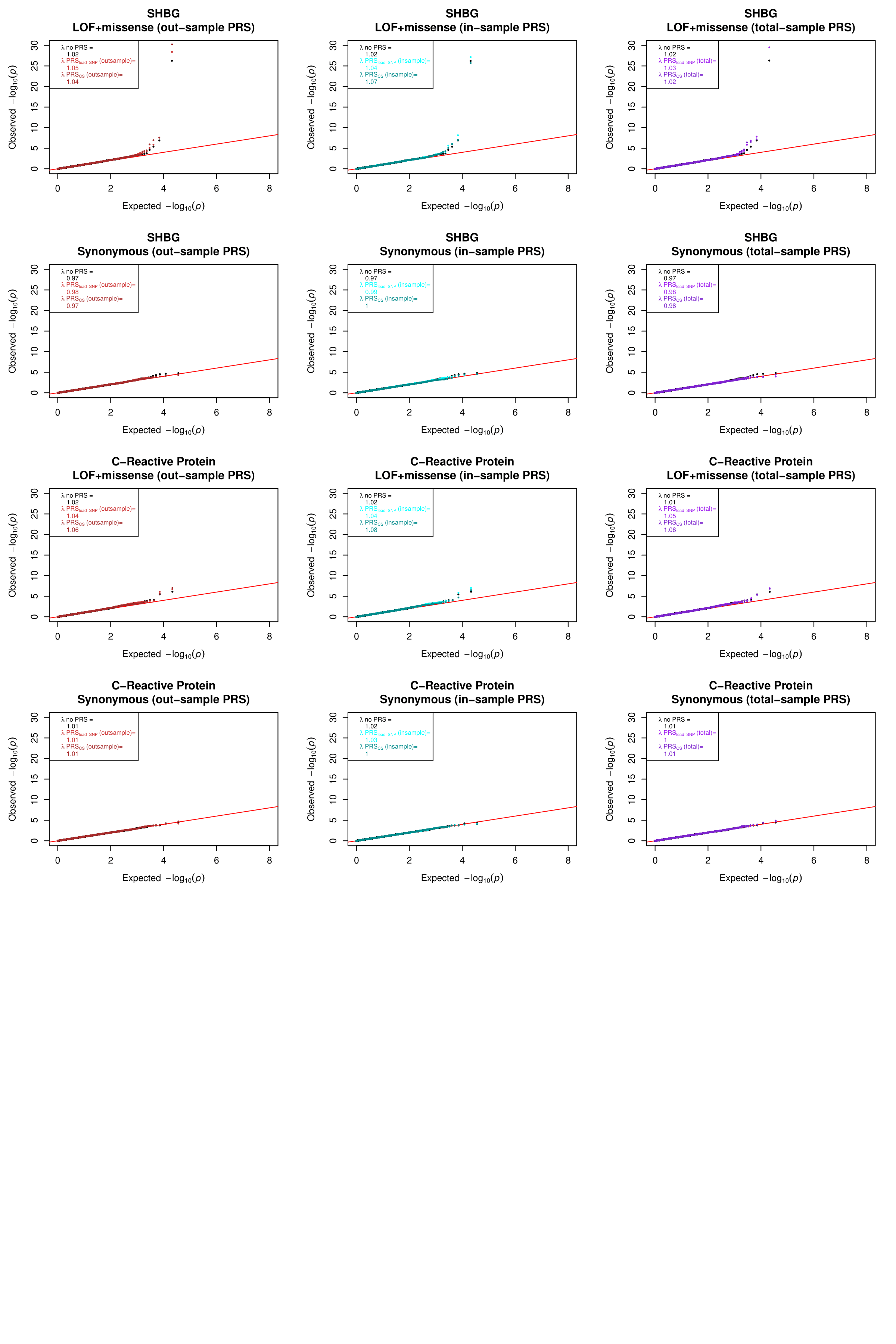
**Supplementary Figure 9: Quantile-quantile plots for exome-wide rare variant analysis of deleterious variants and synonymous variants by model.** The y-axis shows the *P*-values from exome-wide RVAT, while the x-axis shows the *P*-values expected under the null. Black dots show *P*-values from RVAT models without adjusting for PRS, red shows results from models adjusting for PRS derived from out-of-sample GWAS data (left panels), blue shows results from models adjusting for PRS derived from in-sample GWAS data (middle panels), and purple shows results for PRS derived from total GWAS data (right panels). For each phenotype, QQ-plots for deleterious variants (LOF+missense) are plotted above the results for synonymous variants.
